# The gut-brain vagal axis governs mesolimbic dopamine dynamics and reward events

**DOI:** 10.1101/2025.05.12.653303

**Authors:** Oriane Onimus, Faustine Arrivet, Tinaïg Le Borgne, Sylvie Perez, Julien Castel, Anthony Ansoult, Benoit Bertrand, Nejmeh Mashhour, Camille de Almeida, Linh-Chi Bui, Marie Vandecasteele, Serge Luquet, Laurent Venance, Nicolas Heck, Fabio Marti, Giuseppe Gangarossa

**Affiliations:** Université Paris Cité, CNRS, Unité de Biologie Fonctionnelle et Adaptative, F-75013 Paris, France; Sorbonne Université, CNRS, INSERM, Center for Neuroscience Sorbonne University, Institut de Biologie Paris Seine, F-75005 Paris, France; ESPCI Paris, PSL Research University, Brain Plasticity laboratory, CNRS UMR8249, Paris, France; Center for Interdisciplinary Research in Biology (CIRB), Collège de France, CNRS, INSERM, Université PSL, Paris, France; Institut Universitaire de France (IUF)

**Keywords:** Dopamine, Nucleus accumbens, Palatable food, Drugs of abuse, Vagus nerve

## Abstract

Reward-related processes have traditionally been ascribed to neural circuits centered on the dopamine (DA) system. While exteroceptive stimuli, such as food and drugs of abuse, are well-established activators of DA-neuron activity, growing evidence indicates that interoceptive signals also play a critical role in modulating reward. Among these, the gut-brain vagal axis has emerged as a key pathway, yet its precise contribution to mesolimbic DA-dependent signaling, dynamics and behaviors remains poorly defined.

Here, we combine complementary *ex vivo* and *in vivo* approaches across multiple scales to investigate how the gut-brain vagal axis regulates DA dynamics and reward-related behaviors. We show that gut-brain vagal tone is essential for gating mesolimbic DA system activity and functions, modulating DA-dependent molecular and cellular processes, and scaling both food- and drugs-induced reinforcement.

These findings challenge the traditional brain-centric view of reward processing, supporting a more unified and integrated model in which gut-derived and vagus-mediated interoceptive signals are pivotal in intrinsically shaping motivation and reinforcement. By uncovering the influence of gut-brain vagal communication on mesolimbic DA functions, this work offers new insights into the neurobiological mechanisms underlying both adaptive and maladaptive reward processes, with broad implications for eating disorders and addiction.

## Introduction

The regulation of reward-related behaviors relies on the dynamic interplay of neural circuits, with the mesolimbic and nigrostriatal dopamine (DA) systems serving as central regulators. These two systems are primarily composed of dopamine (DA)-neurons located in the midbrain, notably in the ventral tegmental area (VTA), which gives rise to the mesolimbic pathway, while the substantia nigra pars compacta (SNpc) initiates the nigrostriatal pathway. Dopaminoceptive structures, such as the nucleus accumbens (NAc) and the dorsal striatum (DS), collectively integrate a wide range of DA-related stimuli and contribute to scaling appetitive and reward-related behaviors.

Both natural stimuli (*e.g.*, palatable food) and drugs of abuse share some, but not all, mechanistic properties ^1–3^, enabling them to profoundly modulate DA-neurons activity. These exteroceptive (external) signals shape reinforced behaviors by modulating DA-associated molecular, cellular, and circuit-level processes, ultimately influencing the initiation and maintenance of physiological and/or pathological adaptive responses. While the neuronal mechanisms underlying the mobilization and integration of DA signaling have been extensively studied and characterized ^1,4,5^, emerging evidence suggests that brain reward processing and dynamics are also intricately shaped by bodily-born interoceptive (internal) signals ^6–9^. However, the mechanistic underpinnings of interoception in scaling reward events remain largely unknown.

Interoception, *i.e.* the perception of the body’s internal states, plays a crucial role in regulating homeostatic processes ^10^, behavioral adaptations, and emotional and cognitive functions ^11,12^. Beyond merely informing the brain about physiological conditions, interoceptive pathways influence motivation, decision-making, and reinforcement learning ^12,13^. Among these pathways, the vagus nerve serves as a major bridge between peripheral organs and the brain, relaying metabolic, visceral, and immune signals to key neural substrates of adaptive and maladaptive responses ^7,8,14–18^. Among metabolically active peripheral organs, the gut emerges as a central player in coordinating the body-brain tango through a multitude of long-range mechanisms, including hormonal signaling, microbiota-derived metabolites, and both local and gut-brain neuronal connections ^19–22^.

Although the precise mechanisms remain elusive, recent seminal studies have explored the role of gut-vagal afferents in modulating DA-driven food consumption via a polysynaptic gut-to-brain circuit ^6^. However, the precise contribution of the vagus nerve to DA-driven reinforcement behaviors (palatable food and drugs of abuse) and the involvement of the DA mesolimbic (VTA→NAc) system remains poorly explored and understood.

In this study, we leverage multi-scale *ex vivo* and *in vivo* approaches to investigate whether and how the gut-brain vagal axis shapes dopamine (DA) signaling dynamics and reward-related events. We demonstrate that the constitutive/intrinsic activity of the gut-brain vagal axis is essential for gating the activity of the mesolimbic DA system and for scaling cellular and molecular processes associated with both food and drug rewards. Our findings shed light on the interplay between interoceptive processing and mesolimbic functions in shaping reward processes, further challenging the traditional brain-centric perspective on reward and positive reinforcement. Instead, they strongly support a more integrated framework that considers body-brain dynamics in shaping intrinsic motivational processes.

## Material and methods

### Animals

All experimental procedures were approved by the Animal Care Committee of the Université Paris Cité (CEB-22-2019, APAFiS #24407; CEB-38-2021, APAFiS #35447), and carried out following the 2010/63/EU directive. 8–12 weeks old C57BL/6J male mice (Janvier, France) were used and housed in a room maintained at 22 ±1 °C, with a light period from 7h00 to 19h00. Regular chow diet (3.24 kcal/g, reference SAFE® A04, Augy, France) and water were provided *ad libitum* unless otherwise stated. All procedures were designed to minimize animal suffering and reduce the number of animals used.

### Subdiaphragmatic vagotomy (SDV)

Prior to surgery and during 4-5 post-surgery days, animals were provided with *ad libitum* jelly food (DietGel Boost #72-04-5022, Clear H2O). Animals received Buprécare® (buprenorphine 0.3 mg/kg) and Ketofen® (ketoprofen 10 mg/kg) and were anaesthetized with isoflurane (3.5% for induction, 1.5% for maintenance). During surgery the body temperature was maintained at 37 °C using a heated pad. Briefly, using a binocular microscope, the right and left subdiaphragmatic branches of the vagus nerve were carefully isolated along the lower esophagus/stomach and carefully sectioned in vagotomized animals (SDV mice) or left intact in Sham animals. Mice recovered for at least 3–4 weeks before being used for experimental procedures. All experiments took place within the first 2 months following the end of the recovery period (∼12 weeks from surgery). The efficiency of the SDV procedure was evaluated as previously described ^9^.

### Gastric catheter implantation

Following anesthesia, a midline laparotomy was performed to expose the stomach. A small incision was made in the fundus of the stomach to allow for catheter insertion. The catheter tubing was then carefully inserted through this opening. The catheter was tunneled subcutaneously from the abdominal cavity to the interscapular region. A small incision was made between the shoulder blades to exteriorize the catheter. The exposed portion of the catheter was secured with an external metal cap to maintain its integrity and prevent entry of foreign substances. To ensure long-term patency, the catheters were flushed with sterile water 1-2 times per week throughout the experimental period. This regular maintenance helped prevent catheter occlusion. This surgical approach allows for a direct access to the stomach for long-term studies, minimizing stress to the animals and enabling repeated sampling or administration of substances without the need for multiple interventions, bypassing oral ingestions.

### Tracing and viral constructs

Red Retrobeads™ IX were purchased from Lumafluor Inc. and used according to the manufacturer’s instructions. Unless otherwise specified, all AAV recombinant genomes were packaged in serotype 9 capsids. The AAV-PPTA-Cre (6×10^13^ vg/mL) and AAV-PPE-Cre (8×10^13^ vg/mL) contain an expression cassette consisting of the Cre recombinase driven by the promoter of the PPTA (preprotachykinin, substance P) or PPE (preproenkephalin) genes, which are specifically expressed in D1R- and D2R-SPNs, respectively ^23–25^. Each virus was co-injected with an AAV-FLEX-tdTomato in a 1/10.000 ratio to visualize sparse neurons (*ex vivo* electrophysiology and dendritic spine density).

pAAV-FLEX-tdTomato was a gift from Edward Boyden (Addgene viral prep #28306-AAV9; http://n2t.net/addgene:28306 ; RRID:Addgene_28306).

pAAV-hsyn-GRAB_DA2m was a gift from Yulong Li (Addgene viral prep #140553-AAV9; http://n2t.net/addgene:140553; RRID:Addgene_140553). pENN.AAV.CamKII.GCaMP6f.WPRE.SV40 was a gift from James M. Wilson (Addgene viral prep #100834-AAV9; http://n2t.net/addgene:100834 ; RRID:Addgene_100834).

pAAV-hSyn-dLight1.2 was a gift from Lin Tian (Addgene viral prep #111068-AAV5; http://n2t.net/addgene:111068 ; RRID:Addgene_111068).

### Stereotaxic surgery

For all surgical procedures, mice were rapidly anesthetized with isoflurane (3.5%, induction), injected (ip) with the analgesic buprenorphine (Buprecare, 0.3 mg/kg, Recipharm, Lancashire, UK) and ketoprofen (Ketofen, 10 mg/kg, France), and maintained under isoflurane anesthesia (1.5%) throughout the surgery. Mice were placed on a stereotactic frame (David Kopf Instruments, California, USA). Bilateral (AAV-PPTA-Cre + AAV-FLEX-tdTomato or AAV-PPE-Cre + AAV-FLEX-tdTomato) or unilateral (AAV-GRAB^DA2m^) micro-injections were performed at the following coordinates (in mm from bregma): Nac (L= −/+0.9; AP= +1.18, V= −4.4) and DS (L= −/+1.25; AP= +0.95; V=-3.15). For tracing experiments, Red Retrobeads™ IX was injected in the VTA (L= +0.45; AP= −3.4; V=-4.3) for retrograde tracing, whereas AAV9-CamKIIa-GCaMP6f was injected in the PBN (L= +1.5; AP= −5.25; V=-3.35) for anterograde tracing. All viruses were injected at the rate of 0.05 µl/min. Mice recovered for at least 3-4 weeks after the surgery before being involved in experimental procedures.

### *In vivo* fiber photometry

For *in vivo* DA imaging (GRAB^DA2m^, ^26^), a chronically implantable cannula (Doric Lenses, Québec, Canada) composed of a bare optical fiber (400 µm core, 0.48 N.A.) and a fiber ferrule was implanted 100 µm above the location of the viral injection site in the Nac or DS. The fiber was fixed onto the skull using dental cement (Super-Bond C&B, Sun Medical). Real-time fluorescence was recorded using fiber photometry as previously described ^27,28^. Fluorescence was collected in the Nac or DS using a single optical fiber for both delivery of excitation light streams and collection of emitted fluorescence. The fiber photometry setup used 2 light emitting LEDs: 405 nm LED sinusoidally modulated at 330 Hz and a 465 nm LED sinusoidally modulated at 533 Hz (Doric Lenses) merged in a FMC4 MiniCube (Doric Lenses) that combines the 2 wavelengths excitation light streams and separate them from the emission light. The MiniCube was connected to a fiber optic rotary joint (Doric Lenses) connected to the cannula. A RZ5P lock-in digital processor controlled by the Synapse software (Tucker-Davis Technologies, TDT, USA), commanded the voltage signal sent to the emitting LEDs via the LED driver (Doric Lenses). The light power before entering the implanted cannula was measured with a power meter (PM100USB, Thorlabs) before the beginning of each recording session. The light intensity to capture fluorescence emitted by 465 nm excitation was between 25-40 µW, for the 405 nm excitation this was between 10-20 µW at the tip of the fiber. The fluorescence emitted by the GRAB was collected by a femtowatt photoreceiver module (Doric Lenses) through the same fiber patch cord. The signal was then received by the RZ5P processor (TDT). On-line real time demodulation of the fluorescence due to the 405 nm and 465 nm excitations was performed by the Synapse software (TDT). Signals were exported to Python 3.0 and analyzed offline. Data are presented as z-score of ΔF/F.

### Behavioral experiments

#### Time-locked palatable feeding

As previously established ^9^, time-locked and intermittent access to a palatable solution (PS: Intralipid 20% w/v + sucrose 10% w/v) was provided 1-hour/day during 10 consecutive days at ∼11 am. Volume (ml) of consumed palatable solution was measured at the end of the session. Sessions were conducted in cages equipped with an automated online measurement system and locomotor activities were measured using an infrared beam-based activity monitoring system (Phenomaster, TSE Systems GmbH, Germany). Experiments for *in vivo* fiber photometry (GRAB-DA^2m^) were performed after stable food-reward consumption.

#### Operant conditioning

Mice were food-restricted and maintained at 90% of their initial body weight to facilitate learning and performance during the whole operant conditioning. Computer-controlled operant conditioning was conducted in 12 identical conditioning chambers (Phenomaster, TSE Systems GmbH, Bad Homburg, Germany) during the light phase, at the same hour every day until the end of the procedure. Each operant wall had two levers (one active and one inactive) located 3 cm lateral to a central pellet dispenser. The reinforcer was a single 20-mg peanut butter flavoured sucrose tablet (TestDiet, Richmond, USA). Operant training was carried out daily with no interruption for 45 min under a fixed-ratio 1 (FR1, 1 lever press=1 pellet). When the discrimination score between active and inactive lever press (active lever presses/inactive lever presses) exceeded chance level, mice were shifted to sessions under a FR5 (5 lever presses=1 pellet) and a progressive ratio (PR) [3 lever presses more for each subsequent reinforcer (r=3N+3; N=reinforcer number)].

#### Conditioned-place preference (CPP)

The CPP paradigm was performed during the light phase either in food-restricted (maintenance at 90% of initial body weight) for HFD-induced CPP or normally fed mice for wheel running- and psychostimulants-induced CPP. All the compartments were cleaned before each conditioning session. Locomotor activity was recorded with an infrared beam-based activity monitoring system and analyzed with the provided software (Phenomaster, TSE Systems GmbH, Bad Homburg, Germany). The least preferred compartment during the exploration phase was designated as the reward (HFD, drugs of abuse or unblocked wheel)-baited compartment whereas the more preferred compartment as the control (chow, saline or blocked-wheel)-baited compartment. Animals with more than 70% of preference for a compartment on the pre-test day were removed. To reduce anxiety, during the first two days, animals were carefully put in the middle of the apparatus and allowed to freely explore the two compartments for 1h. The subsequent days included alternating conditioning sessions. After 8 days of conditioning [4 sessions in each compartment (chow/saline/blocked wheel *vs* HFD/drugs/unblocked wheel)], for the test day, animals freely explored the two compartments for 20 minutes without cues. The time spent in the reward-paired compartment before *vs* after conditioning was the primary outcome variable.

#### T-Maze

Mice were food-restricted (90 % of initial body weight) during the whole paradigm and tested for positive conditioning in a T-maze apparatus (arm: 35 cm length, 25 cm height, 15 cm width). First, they were habituated to the apparatus (15 min of exploration) for two consecutive days. Then, mice underwent a 5-days training protocol with one arm reinforced with a palatable food pellet (HFD, Research Diets, cat #D12492, 5.24 kcal/g). Each mouse was placed at the starting point and allowed to explore the maze by choosing one of the two arms (reinforced and unreinforced arms). The chosen arm was then blocked for 20 seconds, and the mouse replaced again in the starting arm. This process was repeated for 10 sessions per day.

#### Food preference and choice

Chow *ad-libitum* fed mice were exposed for 1h/day to 3 types of food pellets (HFHS, Research Diets, cat D12451, 4.36 kcal/g; HFD, Research Diets, cat #D12492, 5.24 kcal/g and regular chow diet (3.24 kcal/g, reference SAFE® A04, Augy, France). To avoid food neophobia, mice were pre-exposed to the different diets one week prior to the food-choice test. Mice were weighted and the amount of food eaten during the food choice paradigm was collected at 30min and 1h exposure.

#### Intragastric perfusion coupled to in vivo fiber recordings

Catheters were connected to syringes placed on an infusion pump. According to the experiment, intragastric perfusions were performed at a rate of 100 uL/min for 5 or 10 minutes. A maximum of 1 mL solution was administered in total. For habituation, freely moving mice were perfused with sterile water for 5 minutes at least 2 times before any other experiments. Dopamine dynamics was assessed using fibber photometry with DA biosensor (GRAB-DA2m) *in vivo* (as described above) while perfusing mice. For water perfusions: sterile water was administered for 5 minutes. For high-fat-high sugar perfusions: a solution of 20% intralipids (SIGMA-ALDRICH), and 10% sucrose was administered for 5 or 10 minutes.

#### Drugs-induced locomotor activity

Locomotor activity induced by GBR12909 (10 mg/kg, Tocris, #0421), cocaine (15 mg/kg, Sigma-Aldrich, #C5776), amphetamine (2 mg/kg, Tocris, #2813), morphine (10 mg/kg), or SKF-81297 (5 mg/kg, Tocris, #1447) was recorded in an automated online measurement system using an infrared beam-based activity monitoring system (Phenomaster, TSE Systems GmbH, Bad Homburg, Germany).

#### Haloperidol-induced catalepsy

Animals were injected with haloperidol (0.5 mg/kg, Tocris, #0931) one hour before the catalepsy test. At t=0, 15, 30, 45, 60, 75, 90 minutes, animals were taken out of their home cage and placed in front of a 4 cm elevated steel bar, with the forelegs placed upon the bar while the hind legs remained on the ground surface. The time during which animals remained still was measured. Animals that failed to remain on the bar for at least 30 seconds during the whole test were excluded. A behavioral threshold of 180 seconds was set so the animals remaining in the cataleptic position for this duration were put back in their cage until the next time point.

### Tissue preparation and immunofluorescence

Mice were anaesthetized with pentobarbital (500 mg/kg, Dolethal, Vetoquinol, France) and transcardially perfused with cold (4 °C) PFA 4% for 5 min. Brains were post-fixed in PFA 4% at 4°C for 24h and changed in PBS 1X. 40 μm coronal sections were processed using a vibratome (Leica). Confocal imaging acquisitions were performed after immunohistochemistry protocol, using a confocal microscope (Zeiss LSM 710) as previously described ^29^. The following primary antibodies were used: rabbit anti-Ser235/236-S6 (1:500, Cell Signaling Technology, #2211), guinea pig or rabbit anti-cFos (1:1000, Synaptic Systems, #226 003, 1:1000, Cell Signaling, #2250), mouse anti-TH (1:500 or 1:1000, Millipore, #MAB318). Sections were incubated for 60 min with the following secondary antibodies: goat anti-chicken Alexa488 (1:1000, Invitrogen), donkey anti-rabbit Cy3 AffiniPure (1:1000, Jackson Immunoresearch, 711-165-152), goat anti-mouse Alexa488 (1:500, Invitrogen A21121) or donkey anti-mouse Cy5 (Jackson Immunoresearch, 715-175-150). Structures were selected according to the following coordinates (from bregma, in mm): DS/Nac (1.18 to 0.98), VMH/Arc (−1.46 to −1.82), PBN (−5.02 to −5.34), and AP/cNTS (−7.32 to −7.76). The objectives (10X or 20X) and the pinhole setting (1 airy unit) remained unchanged during the acquisition of a series for all images. Quantification of immunopositive cells was performed using the cell counter plugin of ImageJ taking a fixed threshold of fluorescence as standard reference.

For the imaging and analysis of TH-positive varicosities, image stacks were acquired with Leica SP5 confocal microscope equipped with 1.4 numerical aperture 63x objective with pinhole aperture set to 1 airy unit, pixel size of 60 nm and 200 nm z-step. Image stacks were deconvolved with Huygens software using an experimental psf obtained from 100 nm fluorescent beads. Segmentation of TH-positive varicosities was performed in 3D with the spot segmentation procedure of the ImageJ plugin 3DimageSuite. Image stacks corresponded to a volume of 60×60x3 microns, the number of varicosities were normalized to a cube of 10 microns side.

### *In vivo* electrophysiology on anesthetized animals

Induction of anaesthesia was done with gas mixture of oxygen (1 L/min) and 3% isoflurane (IsoFlo) through a TeamSega apparatus. Mice deeply anesthetized were then placed in a stereotaxic frame (David Kopf), maintained under anesthesia at 2 % isoflurane. Glass electrodes (tip diameters of 1-2 μm, impedances of 6–9 MΩ, filled with 0.5% sodium acetate) were lowered into the VTA (coordinates: 3.1 ± 3 mm posterior to bregma, 0.4 to 0.5 mm lateral to the midline, 3.9 to 5 mm ventral from the brain). Electrical signals were amplified by a high-impedance amplifier (Axon Instruments) and monitored through an audio monitor (A.M. Systems Inc.). The unit activity was digitized at 12.5 kHz and recorded using Spike2 software (Cambridge Electronic Design). Extracellular identification of putative dopamine neurons was based on their location as well as on the set of unique electrophysiological properties that distinguish dopamine from non-dopamine neurons in vivo: (*i*) a typical triphasic action potential with a marked negative deflection; (*ii*) a long duration (>2.0 ms); (*iii*) an action potential width from start to negative trough >1.1 ms; (iv) a slow firing rate (<10 Hz and >1 Hz). Electrophysiological recordings were analyzed using the R software (https://www.r-project.org). DA cell firing was analyzed with respect to the average firing rate and the percentage of spikes within bursts (%SWB, number of spikes within burst divided by total number of spikes). Bursts were identified as discrete events consisting of a sequence of spikes such that: their onset is defined by two consecutive spikes within an interval <80 ms whenever and they terminate with an inter-spike interval >160 ms.

The firing rate and %SWB of each recorded neuron are used as Independent variable. Normality of data set is tested by Shapiro test. For firing frequency and %SWB, comparison of distribution between groups is made using Kolmogorov-Smirnov test. For comparison of the mean value, because the data set exhibits non-normal skewed distribution, comparison between groups is made using a surrogate-based permutation test. For %SWB, we first calculate the absolute difference of mean %SWB between the two groups (Δ<%SWB>o). We then generated 10 000 surrogate data (Δ<%SWB>S). A surrogate data is obtained by resampling two groups from the original dataset (by permutation) and for each resampling by calculating the difference in the mean %SWB of the two resampled groups. Original dataset is obtained by pooling the %SWB observed in Sham and SDV mice and resampling by randomly re-assign these values to two groups. We then calculate 10 000 surrogate values Δ<%SWB>S and count the number of times Δ<%SWB>S ≥ (Δ<%SWB>o. The null hypothesis is that all samples (Sham and SDV) come from the same distribution. The difference reaches statistical significance if the surrogate reproduces the absolute difference of mean %SWB less than 500/100/10 times over 10 000 simulations (*p<0.05, **p<0.01, ***p<0.001). The same method is used to compare the mean difference of firing frequency between groups.

### VTA *ex vivo* electrophysiology: patch clamp recordings

Mice were deeply anesthetized by an intraperitoneal injection of a mix of ketamine (150 mg/kg Virbac 1000) and xylazine (60 mg/kg, Rompun 2%, Elanco). Coronal midbrain sections (250 μm) were sliced with a Compresstome (VF-200, Precisionary Instruments) after intracardial perfusion of cold (4°C) sucrose-based artificial cerebrospinal fluid containing (in mM): 125 NaCl, 2.5 KCl, 1.25 NaH_2_PO_4_, 26 NaHCO_3_, 5.9 MgCl_2_, 25 sucrose, 2.5 glucose, 1 kynurenate (pH 7.2, 325 mOsm). After 8 minutes at 37°C for recovery, slices were transferred into oxygenated artificial cerebrospinal fluid (ACSF) containing (in mM): 125 NaCl, 2.5 KCl, 1.25 NaH_2_PO_4_, 26 NaHCO_3_, 2 CaCl_2_, 1 MgCl_2_, 15 sucrose, 10 glucose (pH 7.2, 325 mOsm) at room temperature for the rest of the day. Slices were individually transferred to a recording chamber continuously perfused at 2 mL/minute with oxygenated ACSF. Patch pipettes (4-6 MΩ) were pulled from thin wall borosilicate glass (G150TF-3, Warner Instruments) with a micropipette puller (P-87, Sutter Instruments Co.). Neurons were visualized using an upright microscope coupled with a Dodt gradient contrast imaging, and illuminated with a white light source (Scientifica). Whole-cell recordings were performed with a patch-clamp amplifier (Axoclamp 200B, Molecular Devices) connected to a Digidata (1550 LowNoise acquisition system, Molecular Devices). Signals were low-pass filtered (Bessel, 2 kHz) and collected at 10 kHz using the data acquisition software pClamp 10.5 (Molecular Devices). VTA location was identified under microscope. Identification of dopaminergic neurons was performed by location and by their electrophysiological properties [width and shape of action potential (AP) and after hyperpolarization (AHP)]. To perform recordings of miniature excitatory postsynaptic currents (mEPSCs), we used a potassium gluconate-based intracellular solution containing (in mM): 135 K-gluconate, 10 HEPES, 0.1 EGTA, 5 KCl, 2 MgCl_2_, 2 ATP-Mg, 0.2 GTP-Na, and biocytin 2 mg/mL (pH adjusted to 7.2). Recordings were conducted in the presence of 500 mM of TTX (tetrodotoxin citrate, HelloBio) to block voltage-gated sodium channels. Comparison of mEPSC distribution between groups is made using Kolmogorov-Smirnov test. Comparisons between means mEPSC amplitude and frequency were performed with parametric Student’s t-test for comparing two groups when parameters followed a normal distribution (Shapiro-Wilk normality test with p > 0.05), or Wilcoxon non-parametric test for non-normal distribution.

### DS and nAc *ex vivo* electrophysiology: patch clamp recordings

Animals were terminally anaesthetized using isofluorane. Sagittal striatal slices (350 μm-thick), containing the DS and Nac were cut using a VT1000S vibratome (VT1000S, Leica Microsystems, Nussloch, Germany) in ice-cold oxygenated solution (ACSF: 125 mM NaCl, 2.5 mM KCl, 25 mM glucose, 25 mM NaHCO_3_, 1.25 mM NaH_2_PO_4_, 2 mM CaCl_2_, 1 mM MgCl_2_, 1 mM pyruvic acid). Slices were then incubated in ACSF at 32–34°C for 60 minutes before returning to room temperature. For whole-cell recordings, borosilicate glass pipettes of 5-7 MΩ resistance were filled with a potassium gluconate-based internal solution consisting of (in mM): 122 K-gluconate, 13 KCl, 10 HEPES, 10 phosphocreatine, 4 Mg-ATP, 0.3 Na-GTP, 0.3 EGTA (adjusted to pH 7.35 with KOH, osmolarity ∼295-300 mOsm). Signals were amplified using with EPC10–2 amplifiers (HEKA Elektronik, Lambrecht, Germany). All recordings were performed at 34°C, using a temperature control system (Bath-controller V, Luigs&Neumann, Ratingen, Germany) and slices were continuously superfused with extracellular solution at a rate of 2 ml/min. Recordings were sampled at 10 kHz, using the Patchmaster v2×32 program (HEKA Elektronik). D1R-SPNs (AAV-PPTA-Cre + AAV-FLEX-tdTomato) and D2R-SPNs (AAV-PPE-Cre + AAV-FLEX-tdTomato) were visualized under direct interference contrast with an upright BX51WI microscope (Olympus, Japan), with a 40x water immersion objective combined with an infra-red filter, a monochrome B/W CCD camera (ORCA-ER, Hamamatsu, Japan) and a compatible system for analysis of images as well as contrast enhancement. Current over voltage (I/V) curves were acquired in current-clamp mode with membrane potentials maintained at −75 mV. Only data from fluorescent (tdTomato) SPNs (AAV-PPTA-Cre *vs* AAV-PPE-Cre) were included in the present study. The active and passive electrophysiological properties of SPNs were calculated according to and consistent with a previous study ^30^.

### Dendritic spine morphology and analysis

Dendrites and spines were either labeled with diolistic technique or via viral expression of tdTomato. The labeling, imaging and analysis workflow has been extensively described previously ^31,32^. For diolistic labeling, 50 mg of tungsten beads (Biorad) was mixed with 3 mg of solid green DiI (3,3’-Dioctadecyloxacarbocyanine Perchlorate, Molecular Probes) dissolved in methylene chloride. DiI-coated beads were coated on the inner surface of a PolyVinylPyrrolidone (Sigma-Adrich) pretreated Teflon tube. Helium gas pressure (150 psi) applied through the genegun eject the beads out of the cartridge onto the brain slice. Beads were delivered through a 3-µm pore-size filter (Isopore polycarbonate, Millipore) to avoid clusters. After labeling, slices were kept in PBS at RT for at least 2 h and mounted in Prolong Gold. Image stacks were acquired with confocal microscope (SP5, Leica) and oil immersion 1.4 numerical aperture 63x objective. 561 nm laser intensity was set so each dendrite occupies the full dynamic range while the gain of the low noise Hybrid detector was kept constant. The pinhole aperture was 1 airy unit and pixel size 60 nm with 200 nm z-step. Iterative deconvolution with experimental psf measured from 100 nm beads was performed using Huygens software. For each striatal neuron, a dendritic segment of 50–70 µm in length and distant from at least 50 µm from the soma or after the first branching point was considered. Dendritic shaft was traced in Neuronstudio software, and dendritic spines detected. Thin and mushroom subtypes are defined by Neuronstudio as spines which 2D head diameter is lower or higher than 350 nm, respectively. 3D coordinates of spines generated by Neuronstudio were imported into ImageJ to segment spine heads with spot segmentation procedure using 3DimageSuite plugin, allowing accurate measurement of spine heads volumes.

### Quantitative RT-PCR

The DS, Nac and nodose ganglia were dissected and snap-frozen using liquid nitrogen. All tissues were kept at −80 °C until RNA extraction. Tissues were homogenized in TRIzol/QIAzol Lysis Reagent (Life Technologies) with 3 mm tungsten carbide beads by using the Tissue Lyser III (QIAGEN, 9003240). Total RNA was extracted using the Rneasy Micro Kit (QIAGEN, 74004). The RNA was quantified by using the NanoDrop 1000 spectrophotometer. 500 ng of mRNA from each sample was used for retrotranscription, performed with the SuperScript®III Reverse Transcriptase (Life Technologies) following the manufacturer’s instructions. Quantitative RT-PCRs were performed in a LightCycler 1.5 detection system (Roche, Meylan France) using the Takyon No Rox SYBR MasterMix dTTP Blue (Eurogentec) in 384-well plates according to the manufacturer’s instruction. All primer sequences used in this study are provided in **Suppl. Table 1**. Relative concentrations were extrapolated from the concentration range for each gene. Concentration values were normalized to the house-keeping gene RPL19.

### Western blotting

The mouse head was cut and immediately immersed in liquid nitrogen for 3 seconds. The brain was then removed and dissected on ice-cold surface, sonicated in 200 µL (DS) and 100 µL (Nac) of 1% SDS supplemented with 0.2% phosphatase inhibitors and 1% protease inhibitors, and boiled for 10 minutes. Aliquots (2.5 µL) of the homogenates were used for protein quantification using a BCA kit (BC Assay Protein Quantitation Kit, Interchim Uptima, Montluçon, France). Equal amounts of proteins (15 µg) supplemented with a Laemmli buffer were loaded onto 10% polyacrylamide gels. Proteins were separated by SDS-PAGE and transferred to PVDF membranes using the Trans-Blot Turbo Transfer System (BIO-RAD). The membranes were immunoblotted with the following antibodies: anti-phospho Tyrosine Hydroxylase Ser40 (1:750, Rabbit, Cell Signaling, #2791), anti-total Tyrosine Hydroxylase (1:1000, Rabbit, Millipore, #AB152), anti-VMAT2 (1:1000, Rabbit, Synaptic Systems, #138302), anti-β-actin (1:5000, Mouse, Sigma-Aldrich, #A1978). Detection was based on HRP-coupled secondary antibody binding using ECL. The secondary antibodies were anti-mouse (1:5000, Cell signaling, #7076S) and anti-rabbit (1:1000, Cell Signaling Technology, #7074). Membranes were imaged using the Amersham Images 680. Quantifications were performed using the ImageJ software.

### Metabolic efficiency analysis

Indirect calorimetry was performed as previously described ^9^. Mice were monitored for whole energy expenditure (EE), O_2_ consumption, CO_2_ production, respiratory exchange rate (RER=VCO_2_/VO_2_, V=volume), fatty acid oxidation and locomotor activity using calorimetric cages (Labmaster, TSE Systems GmbH, Bad Homburg, Germany). Ratio of gases was determined through an indirect open circuit calorimeter. This system monitors O_2_ and CO_2_ at the inlet ports of a tide cage through which a known flow of air is ventilated (0.4 L/min) and regularly compared to a reference empty cage. O_2_ and CO_2_ were recorded every 15 min during the entire experiment. EE was calculated using the Weir equation for respiratory gas exchange measurements. Food intake was measured with sensitive sensors for automated online measurements. Mice were monitored for body weight and composition at the entry and exit of the experiment using an EchoMRI (Whole Body Composition Analyzers, EchoMRI, Houston, USA). Data analysis was performed on Excel XP using extracted raw values of VO_2_, VCO_2_ (ml/h), and EE (kcal/h).

### Quantification of monoamines and their metabolites by RP-HPLC

Monoamines and their metabolites were analyzed by Reversed Phase-High-performance Liquid Chromatography (RP-HPLC) using a Shimadzu system connected to a Waters 2465 electrochemical detector (HPLC-ED, Waters, USA) with a glassy carbon working electrode (0.7 V, 10 nA). The weighted tissues were suspended in an ice-cold solution containing 0.4% ethylenediaminetetraacetic acid and 0.1M perchloric acid, and homogenized for 2 min using a tissue lyser III (Qiagen). After centrifugation, the supernatant was further analyzed with HLPC-ED. Mobile phase [58.5 mM Sodium Acetate (Sigma, St Louis, USA), 0.7 mM Octane Sulfonic Acid (Sigma O 0133, St Louis, USA), pH 3.8 for mobile phase A and 100% MeOH for mobile phase B] was pumped at a flow rate of 1 mL/min and monoamines and metabolite concentrations were detected at an oxidation potential of 750 mV compared to the reference electrode. Compounds were separated by an isocratic flow (86% A / 14% B) using a Phenomenex Chrome-clone 5µm C18 column (length 250 mm; internal diameter 4.6 mm; particle size 5 mm) at 31°C. Monoamines and metabolites were quantified using LabSolution software (Shimadzu, Kyoto, Japan) by integration of the peak absorbance area, employing a calibration curve established with known monoamine concentrations.

### Oral glucose-, insulin-tolerance tests (OGTT and ITT) and insulin dosage

Animals were fasted 6 hours before oral gavage of glucose (3 g/kg) or administration of insulin (0.5 U/kg). Blood glucose was measured from the vein blood tail using a glucometer (Menarini Diagnotics, Rungis, France) at 0, 15, 30, 45, 60, 90, and 120 min. For the OGTT, blood samples were taken at 0, 15, 30 and 60 min, kept on ice until plasma extraction and stock at −80°C until Insulin dosage. Insulin levels were assessed using an insulin Elisa kit, according to the manufacturer’s instructions (mouse ultrasensitive insulin ELISA kit, ALPCO, Salem, USA).

### Re-clustering and transcriptomics meta-analysis

Publicly available transcriptomic data ^33^ were downloaded from Gene Expression Omnibus (https://www.ncbi.nlm.nih.gov/geo/, GSE138651) and analyzed using a Python 3.0 pipeline generated in line with the original publications.

### Statistics

All data are presented as mean ± SEM. Statistical tests were performed with Prism 8 (GraphPad Software, La Jolla, CA, USA). Normality was assessed by the D’Agostino-Pearson test. Depending on the experimental design, data were analyzed using either Student’s t-test with equal variances, one-way ANOVA or two-way ANOVA. The significance threshold was automatically set at p<0.05. ANOVA analyses were followed by Bonferroni post hoc test for specific comparisons only when overall ANOVA revealed a significant difference (at least p<0.05).

## Results

### The gut-brain vagal axis is necessary for food-mediated reward behaviors

We and others have recently highlighted the key role of the gut-brain vagal axis as an active regulator of complex behaviors including cognitive functions ^20,34,35^ and food-driven reward behaviors ^6,7,9,15^. In this line, we have recently shown that peripheral endocannabinoids contribute, at least in part, to reward-driven eating disorders (*i.e.,* binge eating) by acting through the gut-brain vagal axis ^9^. Building on this, here we investigated whether the interoceptive gut-brain vagal axis plays a crucial role in the development of an escalated consumption of food using a chronic and time-locked exposure to a palatable solution (10% sugar and 20% intralipids). To chronically and unbiasedly disrupt the gut-to-brain interoceptive signaling, we employed the subdiaphragmatic vagotomy (SDV) model. While Sham mice exhibited a rapid increase in food consumption across 10 days, SDV mice showed a slower and lower rate of palatable food consumption (**Fig. 1A**). Chronic, time-locked (1h/day) exposure to palatable food is known to induce reward-dependent anticipatory behaviors ^9^, where food consumption relies on a form of positive/reinforced conditioning rather than on metabolic states or needs ^9,36^. Using our model, in combination with telemetric locomotor activity monitoring, we observed that Sham mice displayed an increased locomotor activity before (food-anticipatory activity, FAA) and during (consumption) food intake (**Fig. 1B, C**). In contrast, SDV mice exhibited dampened locomotor activity during both phases (**Fig. 1B, C**). This reduction was not due to pre-existing locomotor deficits, as both Sham and SDV mice had similar locomotor profiles during the dark period (foraging period) (**Fig. 1B**) or in basal conditions (**Suppl. Fig. 1**). In addition, the differences observed in anticipatory and consummatory behaviors were unlikely due to altered metabolic states (**Suppl. Fig. 1**), neither food preferences/palatability, as both groups showed similar preferences and consumption patterns when given a free choice between chow diet (CD), high-fat diet (HFD), and high-fat high-sugar diet (HFHS) (**Fig. 1D**). Given these findings in a time-locked food conditioning paradigm, we next examined the implication of the gut-brain vagal axis in other forms of reward-elicited conditioning. Sham and SDV mice underwent an HFD-induced conditioned-place preference (CPP) paradigm (**Fig. 1E**). Sham mice exhibited a stronger preference for the HFD-baited compartment than SDV mice (**Fig. 1F, F^1^**), despite similar body weight loss (**Fig. 1G**) and food consumption (**Fig. 1H**) during the conditioning protocol, suggesting a role for the vagus nerve in the reinforcing properties of food. To further examine the role of the gut-brain vagal axis in food-conditioned positive responses, we used two additional paradigms: the T-maze and operant lever press paradigms. In the T-maze test (**Fig. 1I**), SDV mice performed worse than Sham mice in associating the HFD-baited arm with the correct choice, demonstrating an impaired reward-driven discrimination (**Fig. 1J**). To assess seeking behaviors, incentive salience, and the motivational properties of food cues, Sham and SDV mice underwent an operant lever press protocol. During the fixed ratio 1 (FR1) schedule, there were no major differences between groups in the number of active lever presses (**Fig. 1K**) and collected rewards (**Fig. 1L**), indicating intact liking and learning abilities. However, under the FR5 schedule, SDV mice displayed a reduced performance (fewer lever presses and collected rewards) over consecutive sessions compared to Sham mice (**Fig. 1K, L**). This deficit was unlikely due to learning impairments, as both groups exhibited similar discrimination between active and inactive levers (**Fig. 1M**) during the conditioning protocol. The reduced performance under FR5 suggested that SDV mice might have lower motivation to work/seek for palatable food. To further explore this food-motivated component, we implemented a progressive ratio (PR) schedule. SDV mice displayed significantly fewer active lever presses and collected rewards than Sham mice (**Fig. 1N, O**), reinforcing the notion of a dampened motivational drive. One potential alternative explanation is that SDV mice exhibit pre-existing metabolic changes that may contribute to their altered food-conditioning responses and motivational drive. However, apart from a reduction in fat mass and a small decrease in energy expenditure during the light phase (**Suppl. Fig. 1A, F**), no significant differences were observed between Sham and SDV mice in body weight, lean mass, food intake, respiratory exchange ratio (RER), fatty acid oxidation (FAO), locomotor activity, or glucose metabolism (**Suppl. Fig. 1**).

**Figure 1:**
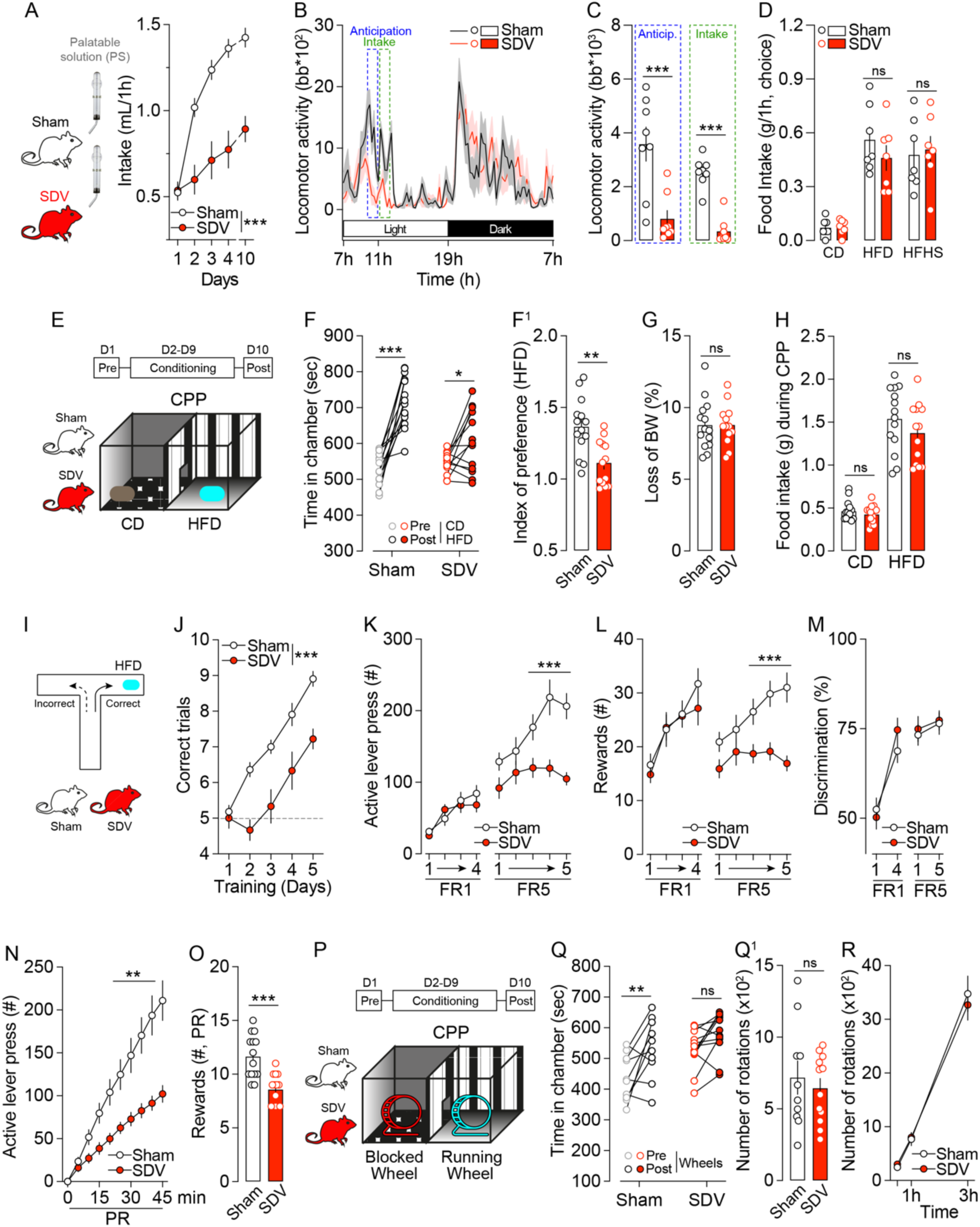
The gut-brain vagal axis is necessary for natural rewards-mediated behaviors. (**A**) Time-locked palatable feeding (binge-like protocol) in Sham (n=8) and SDV (n=8) mice. (**B**) Longitudinal locomotor activity during the binge-like protocol and (**C**) cumulative locomotor activity of Sham (n=8) and SDV (n=8) 1h before (anticipation) and 1h after (intake) access to the palatable solution (PS). (**D**) Food intake during a food preference test with chow diet (CD), high-fat diet (HFD) and high-fat high-sugar diet (HFHS) in Sham (n=7) and SDV (n=7) mice. (**E**) Drawing illustrates the protocol of HFD-induced conditioned-place preference (CPP). (**F**) Time spent in the conditioning chambers and (**F^1^**) index of preference following the CPP protocol for Sham (n=14) and SDV (n=13) mice. (**G**, **H**) Loss of body weight (BW) and HFD intake during the CPP protocol. (**I**) Drawing illustrates the protocol of T-Maze test. (**J**) Number of correct trials (HFD-baited arm) in Sham (n=11) and SDV (n=9) mice. Number of active lever presses (**K**), rewards collected (**L**) and active/inactive lever discrimination (**M**) during the FR1 and FR5 schedules of the operant conditioning in Sham (n=14) and SDV (n=14) mice. Number of active lever presses (**N**) and rewards collected (**O**) during the PR schedule of the operant conditioning in Sham (n=14) and SDV (n=14) mice. (**P**) Drawing illustrates the protocol of running wheel-induced CPP. (**Q**) Time spent in the conditioning chambers and (**Q**^1^) number of wheel rotations during conditioning for Sham (n=10) and SDV (n=11) mice. Statistics: *p<0.05, **p<0.01 and ***p<0.001 for specific comparisons. Two-way ANOVA (A, C, D, F, H, J, K, L, M, N, Q, R), Student’s t-test (F^1^, G, O, Q^1^).

To extend our findings to another food-independent natural reward (*i.e.,* physical activity), we conducted a CPP test using a running wheel paradigm (**Fig. 1P**). While Sham mice displayed a positive conditioning (*e.g.,* preference for the wheel running compartment), SDV mice did not (**Fig. 1Q**), despite comparable wheel rotations during conditioning sessions (**Fig. 1Q^1^**) and similar endurance in a three-hour running wheel test (**Fig. 1R**).

Altogether these results indicate that the gut-brain vagus nerve is necessary for allowing the establishment of natural rewards-induced positive reinforcement and conditioning.

### The gut-brain vagal axis is essential for reward behaviors driven by drugs of abuse

Given that also non-food natural rewards depended on the gut-brain vagal axis and since natural rewards and drugs of abuse share similar properties in driving positive reinforcement and conditioning, we investigated the role of the gut-brain vagal axis in mediating the psychoactive (locomotor activity, **Fig. 2A**) and rewarding properties (CPP) of drugs of abuse. As these substances directly activate the mesolimbic reward system by promoting the release of dopamine (DA), this strategy allows us to explore the potential of the vagal pathway in regulating the complex mechanisms of reward-related processes independently of food-related stimuli. Thus, Sham and SDV mice were administered with cocaine (15 mg/kg), a psychostimulant which, by blocking the DA transporter (DAT), promotes the accumulation of DA and triggers an increase in locomotor activity. Astonishingly, SDV mice showed a lower cocaine-induced locomotor activity during both acute and chronic (sensitization phase) administration (**Fig. 2B, C**). As the response to cocaine cannot only be attributed to DA, we evaluated DA-mediated responses by using a specific DAT blocker. Therefore, mice were administered with GBR (10 mg/kg), with SDV mice showing a significant lower GBR-induced locomotor activity than Sham controls (**Fig. 2D, D^1^**). However, when animals were administered with amphetamine (2 mg/kg), a psychostimulant which, beyond its similar actions with cocaine, actively releases DA, no differences were observed (acute and chronic exposure) (**Fig. 2E**). Next, we used morphine (10 mg/kg), a drug of abuse which promotes DA release by disinhibiting VTA DA-neurons ^37,38^, and again observed a reduction in the elicited locomotor response in SDV mice (**Fig. 2F, F^1^**). These results suggest that the vagus nerve may modulate DA dynamics and/or its postsynaptic integration.

**Figure 2:**
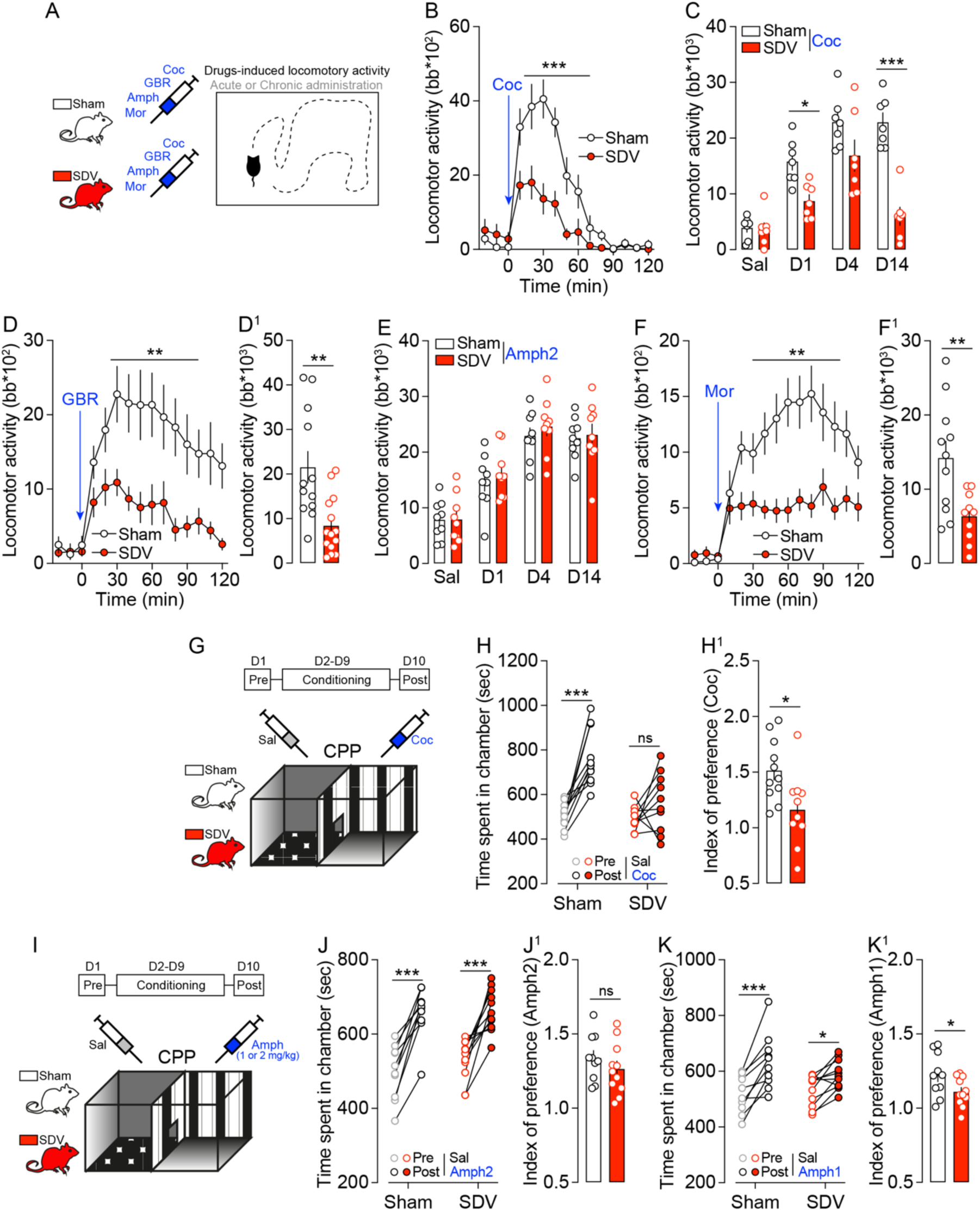
The gut-brain vagal axis is essential for drugs of abuse-mediated reward behaviors. (**A**) Drawing illustrates the protocol of drugs-induced locomotor activity. Acute (**B**) and repeated (**C**, cumulative responses) locomotor activity induced by cocaine (15 mg/kg) in Sham (n=7) and SDV (n=9) mice. (**D, D^1^**) Longitudinal and cumulative locomotor activity induced by GBR12909 (10 mg/kg) in Sham (n=12) and SDV (n=14) mice. (**E**) Acute and repeated (cumulative responses) locomotor activity induced by amphetamine (2 mg/kg) in Sham (n=9) and SDV (n=9) mice. (F, F^1^) Longitudinal and cumulative locomotor activity induced by Morphine (10 mg/kg) in Sham (n=11) and SDV (n=10) mice. (**G**) Drawing illustrates the protocol of cocaine-induced CPP. (**H**) Time spent in the conditioning chambers and (**H^1^**) index of preference following the CPP protocol for Sham (n=11) and SDV (n=10) mice. (**I**) Drawing illustrates the protocol of amphetamine-induced CPP. (**J**) Time spent in the conditioning chambers and (**J^1^**) index of preference following the Amph (2 mg/kg)-CPP protocol for Sham (n=10) and SDV (n=11) mice. (**K**) Time spent in the conditioning chambers and (**K^1^**) index of preference following the Amph (1 mg/kg)-CPP protocol for Sham (n=10) and SDV (n=11) mice. Statistics: *p<0.05, **p<0.01 and ***p<0.001 for specific comparisons. Two-way ANOVA (B, C, D, E, F, H, J, K), Student’s t-test (D^1^, F^1^, H^1^, J^1^, K^1^).

However, while these results point to a potential effect of gut interoception on DA-dependent behaviors, they do not reveal yet whether the rewarding and conditioning properties of psychostimulants depend on the integrity of the gut-brain vagal axis. To tackle this key question, we performed a cocaine- and an amphetamine-induced CPP tests (**Fig. 2G, I**). While Sham mice were positively conditioned to cocaine, no significant preference was observed in SDV mice (**Fig. 2H, H^1^**). Interestingly, the effect of amphetamine-induced CPP in SDV mice depended on the conditioning doses. At 2 mg/kg we observed that both experimental groups were positively conditioned (**Fig. 2J, J^1^**). However, when mice were conditioned with a lower dose of amphetamine (1 mg/kg), we observed a lower conditioning index in SDV mice compared to controls (**Fig. 2K, K^1^**), suggesting that the physiological consequences of neuronal adaptations observed in SDV mice may be overridden at higher DA levels. First, these findings reveal that the constitutive/intrinsic activity of the gut-brain vagus nerve plays a crucial role in mediating food- and drugs-reinforced behaviors (**Fig. 1, 2**). Second, they also indicate that the integrity of this interoceptive axis is not only essential for the rewarding properties of psychostimulants that passively increase DA levels by promoting accumulation, but also for those that actively stimulate DA release at terminals. This highlights a potential role for the vagus nerve in regulating the tonic activity of VTA DA-neurons and/or DA integration in the NAc.

### The gut-brain vagal axis is necessary for food-driven DA dynamics

To assess whether and how *in vivo* DA dynamics depend on the integrity of the interoceptive gut-brain vagal axis, we performed *in vivo* recordings of DA transients/fluctuations using fiber photometry in mice intermittently exposed to a palatable solution (PS). Sham and SDV mice were injected with an AAV-GRAB^DA2m^ in the NAc (**Fig. 3A**), and DA dynamics were recorded during food consumption (**Fig. 3A-C**). Overall, in both groups we observed comparable increased DA levels upon palatable food consumption (**Fig. 3B**). However, when analyzing the rapid events occurring around food consumption, we noticed an early rise in NAc DA levels in Sham but not SDV mice before food consumption (**Fig. 3C, C^1^**), most likely corresponding to the food anticipatory phase (**Fig. 3C, C^1^**). Moreover, a significant difference between Sham and SDV mice was also observed during the first seconds of food consumption (**Fig. 3C, C^1^**).

**Figure 3:**
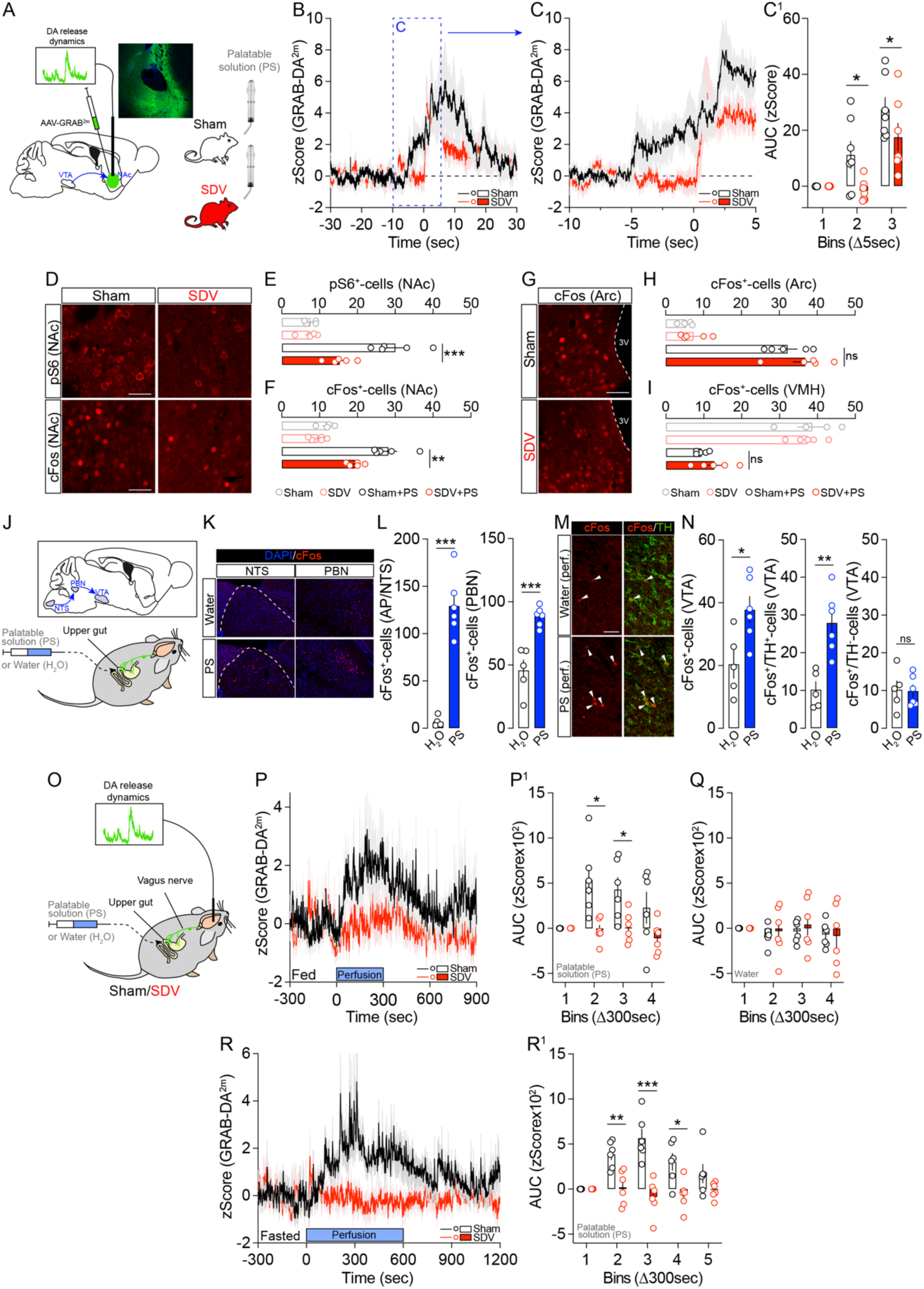
The gut-brain vagal axis is necessary for food-driven DA resealing and molecular dynamics. (**A**) Drawing illustrates our viral strategy to measure *in vivo* DA events (fiber photometry coupled to the DA biosensor GRAB-DA^2m^) in the NAc of Sham and SDV mice during the access to a palatable solution. (**B**) *In vivo* DA dynamics before and during food access. (**C, C^1^**) Inset shows rapid DA transients at a lower time scale and quantified (zScore AUC) periods (1 bin = 5 sec) in Sham (n=7) and SDV (n=7) mice. (**D-F**) Immunofluorescence detection of pS6 and cFos and their respective quantifications (pS6, **E**; cFos, **F**) in the NAc of Sham (n=4), SDV (n=5), Sham+PS (n=5) and SDV+PS (n=5) mice. (**G-I**) Immunofluorescence detection of cFos and its quantifications in the hypothalamus (Arc, **H**; VMH, **I**) of Sham (n=4), SDV (n=5), Sham+PS (n=5) and SDV+PS (n=5) mice. (**J**) Drawing illustrates the intragastric perfusion strategy to observe molecular activations in the brainstem and VTA. (**K, L**) Immunofluorescence detection and quantification of cFos in the NTS and PBN following intragastric perfusion of water (n=5) or PS (n=6). (**M, N**) Immunofluorescence detection and quantification of cFos in the VTA following intragastric perfusion of water (n=5) or PS (n=6). (**O**) Drawing illustrates the intragastric perfusion strategy to measure *in vivo* NAc DA transients by fiber photometry. (**P, P^1^**) *In vivo* DA dynamics during intragastric perfusion of PS in fed Sham (n=6) and SDV (n=6) mice. (**Q**) *In vivo* DA dynamics during intragastric perfusion of water in Sham (n=6) and SDV (n=6) mice. (**R, R^1^**) *In vivo* DA dynamics during intragastric perfusion of PS in fasted Sham (n=6) and SDV (n=6) mice. Scale bars: 50 μm. Statistics: *p<0.05, **p<0.01 and ***p<0.001 for specific comparisons. Two-way ANOVA (C^1^, O^1^, P, Q^1^), One-way ANOVA (E, F, H, I), Student’s t-test (K, M).

Intermittent exposure to palatable food also triggers DA-dependent molecular adaptations in the NAc ^9^. Using two proxies of NAc molecular activity (pS6 and cFos, ^39^), we observed a reduction in pS6- and cFos-activated NAc-neurons in SDV mice compared to controls (**Fig. 3D, E, F**). However, no differences of cFos expression profiles were observed between Sham and SDV mice following palatable food consumption in the hypothalamus with similar food-induced cFos activation (arcuate nucleus, Arc) or cFos inhibition (ventromedial hypothalamus, VMH) in both groups (**Fig. 3G, H, I**). These results indicate that the gut-brain vagal axis is necessary for mediating molecular and cellular reward events within the mesolimbic system, while leaving key homeostatic regulatory nodes unaffected.

Food intake mobilizes multiple physiological processes simultaneously. To exclude orosensory inputs from potential gut-to-brain nutritional/metabolic signals, we intragastrically perfused the same amount of palatable solution or water (1 mL, 100 μL/min) in mice implanted with an intragastric catheter (**Fig. 3J**). As expected, perfusion of the palatable solution increased cFos levels in the NTS, the main sensory vagal direct relay, and in the PBN, a key NTS-projecting structure (**Fig. 3K, L**). Interestingly, we also observed an increase in cFos-positive cells in the VTA, notably in TH-positive DA-neurons (**Fig. 3M, N**), with no major changes in TH-negative neurons. Previous studies have shown that PBN-neurons project to the midbrain, including the SNpc ^6^ and VTA ^40,41^. Using retrograde and anterograde viral tracing strategies, we confirmed such PBN-to-VTA connections (**Suppl. Fig. 2A, B**), whereas only a few and scattered NTS-projecting neurons were directly connected to the VTA (**Suppl. Fig. 2A, B**).

To fully assess vagus-dependent *in vivo* VTA-to-NAc DA dynamics during intragastric perfusion of palatable food, Sham and SDV mice received an injection of AAV-GRAB^DA2m^ in the NAc and were subsequently implanted with an intragastric catheter (**Fig. 3O**). While intragastric perfusion of the palatable solution in fed Sham mice led to an immediate increase in NAc DA levels (**Fig. 3P, P^1^**), no changes in DA dynamics were observed in fed SDV mice (**Fig. 3P, P^1^**). Since the vagus nerve also conveys mechanical information ^42^, both Sham and SDV mice were perfused with an equal volume of water. No modulation of NAc DA events was recorded in both groups (**Fig. 3Q**), indicating that mechanical forces do not impact *in vivo* DA dynamics. Lastly, we investigated whether the absence of DA release events in SDV mice depended on the metabolic state (fed mice in **Fig. 3P, P^1^**). Fasted mice were intragastrically perfused with a palatable solution for a longer period (10 min), and again, no NAc DA releasing transients were observed in SDV mice as compared to Sham controls (**Fig. 3R, R^1^**).

These results reveal that the gut-brain vagal axis represents a key route of interoceptive information, capable of modulating mesolimbic, and not only nigrostriatal ^6,15^, food-dependent DA dynamics.

### The gut-brain vagal axis is necessary for drugs-associated DA dynamics

Next, we investigated whether the integrity of the gut-brain vagal axis was necessary for mediating drug-associated *in vivo* DA dynamics, which may explain, at least in part, the behavioral effects observed with drugs of abuse (**Fig. 2**). To assess this and measure psychostimulants-evoked DA release, Sham and SDV mice were injected with AAV-GRAB^DA2m^ in the NAc (**Fig. 4A**) or DS (**Fig. 4B**). Surprisingly, while cocaine strongly increased NAc DA levels, SDV mice exhibited significantly reduced cocaine-evoked DA accumulation (**Fig. 4C, C^1^**). This finding was corroborated using AAV-dLight1.2, another DA biosensor (**Suppl. Fig. 3**). However, when DA dynamics were measured in the DS, no differences in cocaine-evoked DA accumulation were observed between groups (**Fig. 4D, D^1^**). Then, we examined *in vivo* DA dynamics following amphetamine administration. While amphetamine (2 mg/kg) triggered similar DA release/accumulation at later time points (no differences during the last 10 min of recording) (**Fig. 4E, E^1^**), SDV mice displayed a noticeably slower induction of NAc DA release/accumulation during the first 10 min post-administration (**Fig. 4E, E^1^**). As with cocaine (**Fig. 4D, D^1^**), no differences in amphetamine-evoked DA release/accumulation were observed in the DS of both experimental groups (**Fig. 4F, F^1^**). Since DA modulates the molecular activity of NAc-neurons ^43^ (see also **Fig. 3D-F**), we examined postsynaptic molecular adaptations within the mesolimbic DA system. Compared to Sham mice, SDV mice exhibited a reduced cocaine-induced pS6 and cFos activation in the NAc (**Fig. 4G, H**). One possible explanation is that psychostimulants might directly act on the vagus nerve, therefore leading to the dampened effects observed in SDV mice. To investigate this hypothesis, we first performed a single-cell transcriptomic meta-analysis ^33^ of vagal nodose ganglia (*Phox2b*- and *Slc17a6*-enriched neurons) and found no co-expression of Slc6a3 (DAT transcript) (**Fig. 4I**), suggesting that vagal neurons do not directly express the DA transporter which is necessary for the action of cocaine and amphetamine. Next, we examined molecular responses in the nodose ganglia following cocaine administration. While cocaine increased *cFos* and *Arc* mRNA levels in the NAc (**Fig. 4J**), no changes in the expression of these key immediate early genes were detected in the nodose ganglia of cocaine-treated mice (**Fig. 4K**). These results reveal that the gut-brain vagal axis is essential for scaling the physiological activity of the mesolimbic DA system, and that the vagal integrity is crucial for reward-related events.

**Figure 4:**
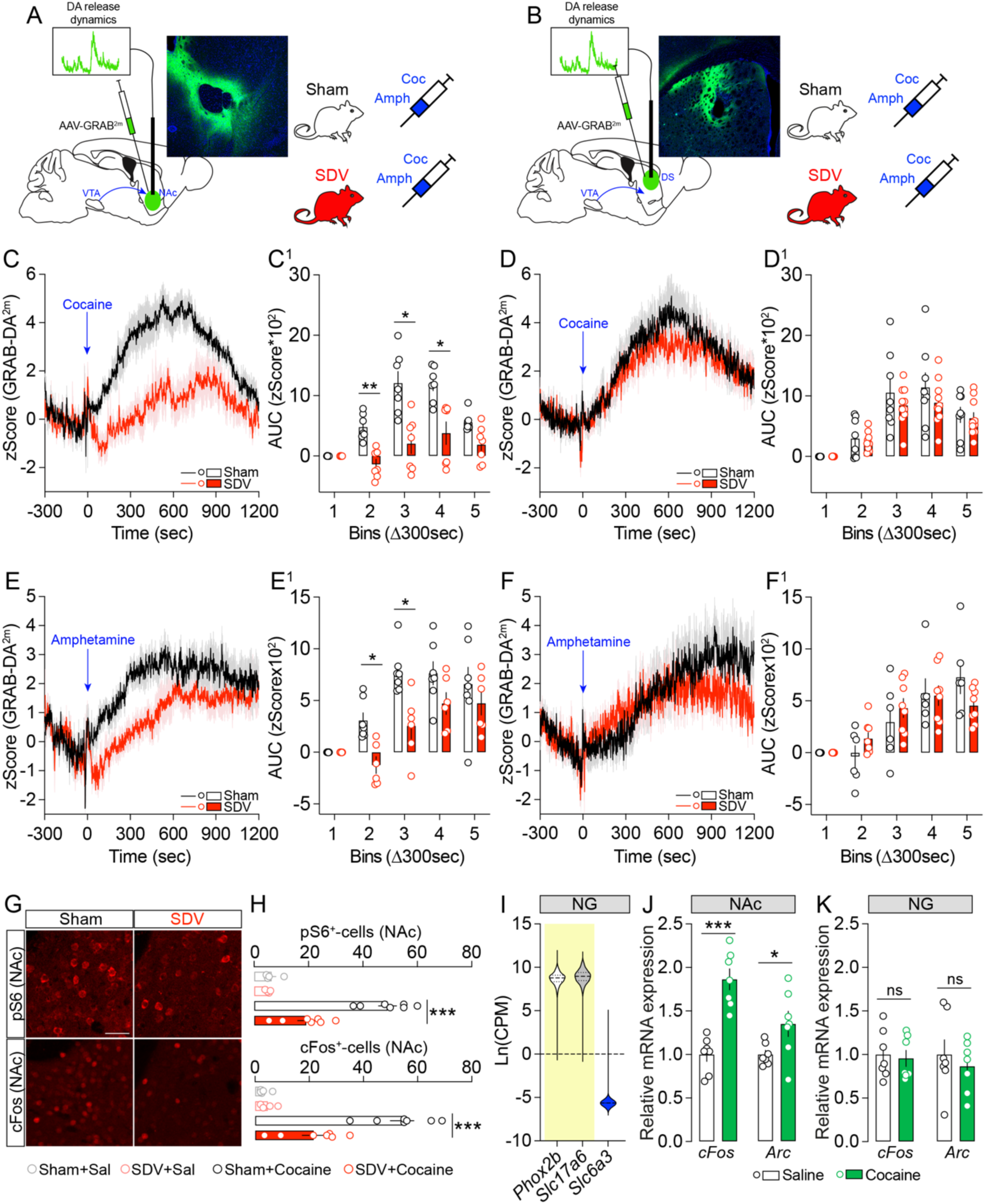
The gut-brain vagal tone plays a permissive role in drugs-induced DA resealing and molecular dynamics. (**A, B**) Drawings illustrate the viral strategy used to measure psychostimulants-induced DA dynamics in the NAc (**A**) and the DS (**B**). (**C, C^1^**) *In vivo* DA dynamics during cocaine-induced DA release/accumulation in the NAc of Sham (n=7) and SDV (n=7) mice. (**D, D^1^**) *In vivo* DA dynamics during cocaine-induced DA release/accumulation in the DS of Sham (n=8) and SDV (n=10) mice. (**E, E^1^**) *In vivo* DA dynamics during amphetamine-induced DA release/accumulation in the NAc of Sham (n=6) and SDV (n=6) mice. (**F, F^1^**) *In vivo* DA dynamics during amphetamine-induced DA release/accumulation in the DS of Sham (n=6) and SDV (n=8) mice. (**G**) Immunofluorescence detection of pS6 and cFos and their respective quantifications (**H**) in the NAc of Sham (n=4), SDV (n=4), Sham+Cocaine (n=7) and SDV+Cocaine (n=7) mice. (**I**) Transcriptomic meta-analysis of *Phox2b*, *Slc17a6* and *Slc6a3* in vagal sensory neurons (n=275). (**J, K**) Expression of cFos and Arc in the NAc (**J**) or nodose ganglia (NG, **K**) of animals treated with Saline (n=7) or Cocaine (n=7). Scale bars: 50 μm. Statistics: *p<0.05, **p<0.01 and ***p<0.001 for specific comparisons. Two-way ANOVA (C^1^, D^1^, E^1^, F^1^), One-way ANOVA (H), Student’s t-test (J, K).

### The integrity of the gut-brain vagal axis is necessary for the spontaneous activity of VTA DA-neurons

To determine whether the gut-brain vagus nerve spontaneously contributes to regulating the activity of VTA DA-neurons, we conducted a series of *in vivo* and *ex vivo* electrophysiological experiments. *In vivo* juxtacellular recordings revealed a blunted VTA DA-neuronal activity in SDV mice (**Fig. 5A**), characterized by a reduced spike frequency and a lower percentage of spikes within bursts (SWB%) (**Fig. 5B, C**). These findings suggest that the integrity of this interoceptive vagal relay is essential for establishing the firing pattern of VTA DA-neurons. To further test this intrinsic dependency of DA-neuron activity on the integrity of the gut-brain vagal axis, we decided to trigger NAc DA releasing events independently of food- or drugs-related stimuli by using the tail suspension (TS) test which promptly activate VTA DA-neurons ^9,44^. Our results indicate that lower TS-induced NAc DA releasing events in SDV mice compared to controls (**Suppl. Fig. 4A, A^1^**).

**Figure 5:**
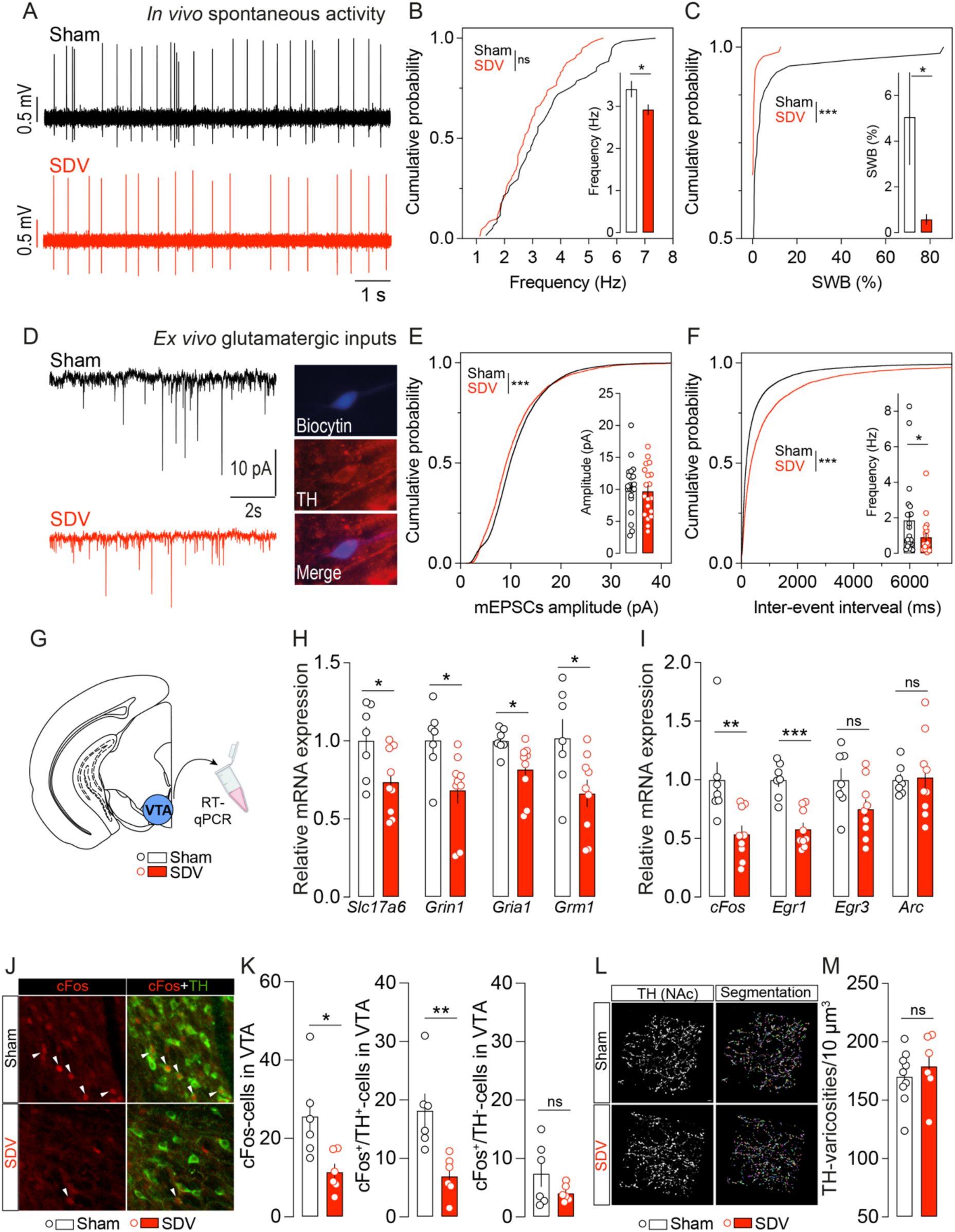
The gut-brain vagal tone regulates the activity of VTA DA-neurons. (**A**) Representative traces of *in vivo* juxtacellular recordings of VTA DA-neurons in Sham and SDV mice. (**B, C**) Spike frequency and percentage of spikes within bursts (SWB%) of VTA DA-neurons in Sham (61 neurons) and SDV (81 neurons) mice. (**D**) Representative traces of *ex vivo* recording to measure the excitatory/glutamatergic drive onto VTA DA-neurons in Sham and SDV mice. Immunofluorescence images show a biocytin-filled VTA DA-neuron. (**E, F**) Amplitude and frequency of mEPSCs onto VTA DA-neurons in Sham (22 neurons) and SDV (19 neurons) mice. (G) Drawing illustrates the area of micro-punch to dissect the mouse VTA for RT-qPCR studies. (**H**) Expression of *Slc17a6*, *Grin1*, *Gria1*, *Grm1* and (**I**) *cFos*, *Egr1*, *Egr3*, *Arc* in Sham (n=7) and SDV (n=9) mice. (**J, K**) Immunofluorescence detection and quantifications of cFos (red) and TH (green) in Sham (n=6) and SDV (n=6) mice. (L, M) 3D reconstruction and computational segmentation of VTA-projecting TH-positive varicosities in the NAc of Sham (n=9) and SDV (n=6) mice. Statistics: *p<0.05, **p<0.01 and ***p<0.001 for specific comparisons. Student’s t-test (E, H, I, K, M), Wilcoxon test (F), Kolmogorov-Smirnov test (B, C, E, F), surrogate-based permutation test (B, C).

Since the excitatory glutamatergic drive onto VTA DA-neurons plays a critical role in determining their firing rate ^37,45–47^, we examined excitatory synaptic inputs in *ex vivo* VTA preparations (**Fig. 5D**). In SDV mice, we observed reduced excitatory input strength, as indicated by a lower cumulative probability of amplitude (though not the average amplitude) (**Fig. 5E**), as well as decreased frequency (both cumulative probability and average frequency) (**Fig. 5F**). To complement these findings, we analyzed the expression of key transcripts involved in glutamatergic transmission efficiency and VTA molecular activity/plasticity (**Fig. 5G**). Consistent with the observed reduction in VTA DA-neuron firing and excitatory drive, SDV mice exhibited lower mRNA expression levels of *Slc17a6*, and genes encoding for glutamate receptors (*Grin1*, *Gria1*, and *Grm1*) (**Fig. 5H**), along with decreased expression of immediate early genes such as *cFos* and *Egr1* (**Fig. 5I**). To further explore this, we assessed the basal expression of cFos (protein) in the VTA of Sham and SDV mice (**Fig. 5J**). SDV mice showed a lower number of cFos-positive cells in the VTA, particularly within VTA DA-neurons (cFos^+^/TH^+^-cells) (**Fig. 5K**).

Given the reduction in reward stimulus-evoked DA release, both in response to food (**Fig. 3**) and psychostimulants (**Fig. 4**), as well as the blunted activity of VTA DA-neurons (**Fig. 5A-K**) in SDV mice, we investigated whether these effects were associated with structural and/or biochemical alterations in DA processes. No significant differences were observed in the NAc and DS between Sham and SDV mice regarding: (*i*) the number of TH-positive varicosities (**Fig. 5L, M** and **Suppl. Fig. 5A**), (*ii*) TH and VMAT2 protein levels (**Suppl. Fig. 5B-D**), (*iii*) monoamines levels (DA, NA, 5-HT) and their metabolites (DOPAC, VMA, HVA, NMN, 3-MT, HIAA) (**Suppl. Fig. 5F, G**), or (*iv*) DA-related metabolic enzyme levels (*Maoa*, *Maob*, *Comt*) (**Suppl. Fig. 5H, I**).

These findings indicate that the gut-brain vagal axis constitutively contributes to regulating and scaling VTA DA-neuronal activity while leaving the basal DA synthetic pathway unaltered (including DA production, degradation and storage).

### The integrity of the gut-brain vagal axis is necessary for the cell type-specific functions of NAc dopaminoceptive neurons

Vagus nerve-mediated changes of the activity of VTA DA-neurons may induce long-lasting adaptations in dopaminoceptive spiny projection neurons (SPNs) of the NAc, potentially contributing to the downregulated behavioral reward-responses observed in SDV mice (**Fig. 1, 2**). Thus, we decided to determine whether the integrity of the gut-brain axis was essential for dopamine D1R- and D2R-dependent behavioral, cellular, and molecular processes. Thus, we pharmacologically challenged Sham and SDV mice using well-characterized D1R- and D2R-specific ligands ^48–51^, notably SKF-81297 (a D1R agonist, 5 mg/kg) and haloperidol (a D2R antagonist, 0.5 mg/kg) (**Fig. 6A**). As expected, SKF-81297 induced a robust increase in locomotor activity in Sham mice (**Fig. 6B**). However, this response was significantly attenuated in SDV mice (**Fig. 6B**). Consistently, SKF-induced cFos activation in the NAc was lower in SDV mice compared to Sham controls (**Fig. 6C**). Next, we administered haloperidol and observed a reduced cataleptic response in SDV mice (**Fig. 6D**). Given that D2Rs are expressed both postsynaptically on D2R-SPNs and presynaptically on DA-neurons/terminals within the mesolimbic reward circuit, we assessed cFos expression as a postsynaptic marker ^50,52^ and TH^S40^ phosphorylation as a presynaptic marker ^53^ in the NAc. Interestingly, SDV mice exhibited a decreased number of haloperidol-induced cFos-positive cells (**Fig. 6E**), suggesting impaired post-synaptic responses. However, TH^S40^ phosphorylation remained unchanged between experimental groups (**Fig. 6F**), indicating that presynaptic DA-mediated D2R signaling was unaffected. These findings suggest that the gut-brain vagal tone is crucial for modulating D1R- and D2R-related behavioral and cellular adaptations.

**Figure 6:**
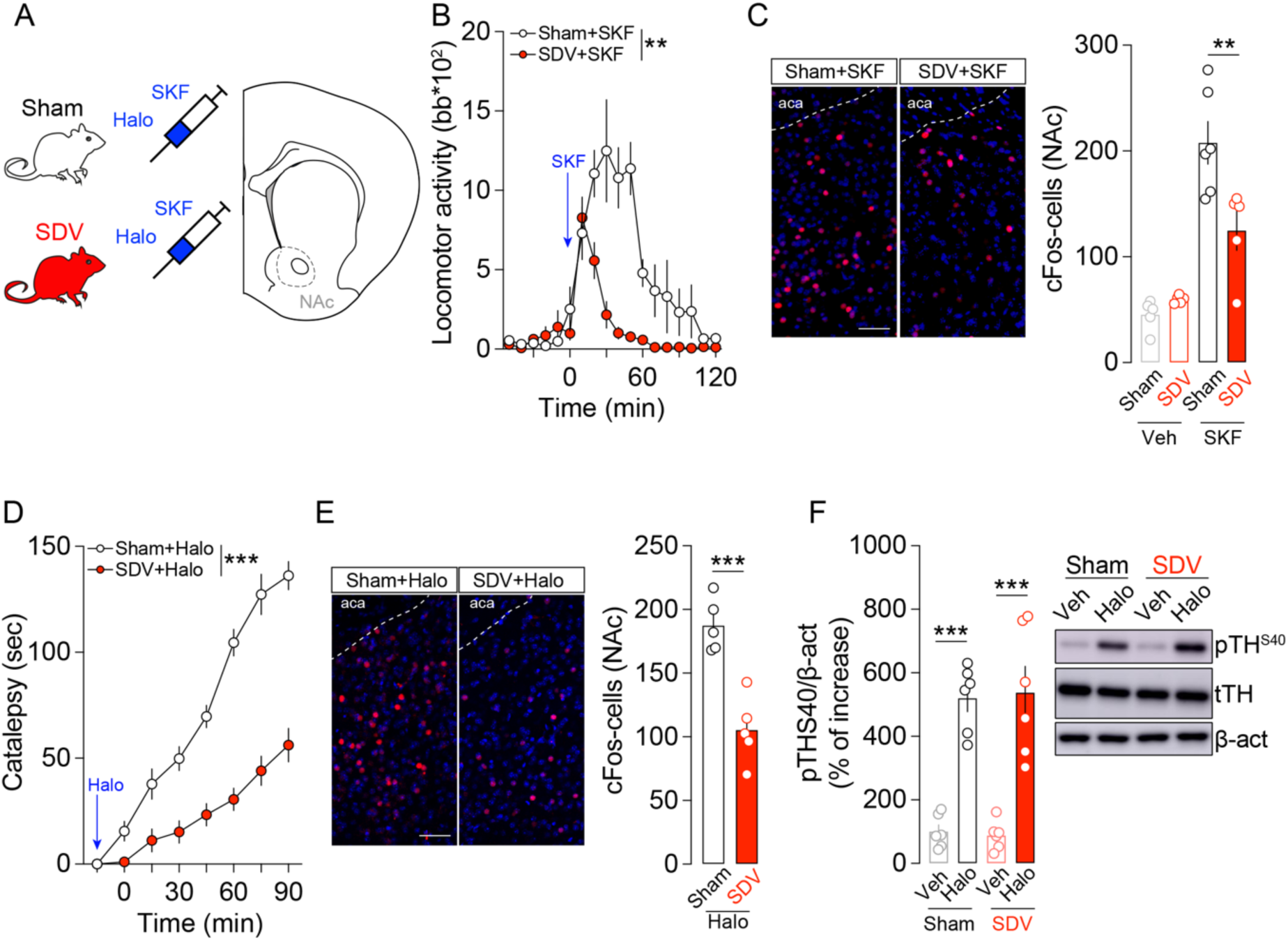
Gut-brain vagal inputs tune D1R- and D2R-dependent processes. (**A**) Drawing illustrates the pharmacological strategy used to evaluate D1R- and D2R-dependent behavioral and signaling events. (**B**) Locomotor activity induced by the D1R agonist SKF81297 (5 mg/kg) in Sham (n=7) and SDV (n=8) mice. (**C**) Immunofluorescence detection and quantification of cFos mice treated with vehicle (Veh) or SKF81297 (SKF): Sham+Veh (n=5), SDV+Veh (n=5), Sham+SKF (n=6), SDV+SKF (n=5). (**D**) Cataleptic response to haloperidol (Halo, 0.5 mg/kg) in Sham (n=7) and SDV (n=7) mice. (**E**) Immunofluorescence detection and quantification of cFos mice treated with haloperidol (Halo) in Sham (n=5) and SDV (n=5) mice. (**F**) Quantifications and representative blots of phosphorylated levels of TH at Ser40 (pTH^S40^) in animals treated with vehicle or haloperidol: Sham+Veh (n=6), SDV+Veh (n=6), Sham+Halo (n=6), SDV+Halo (n=6). Scale bars: 50 μm. Statistics: **p<0.01 and ***p<0.001 for specific comparisons. Two-way ANOVA (B, D), One-way ANOVA (C, F), Student’s t-test (E).

To assess whether D1R- and D2R-SPNs undergo specific adaptive changes, we employed a viral approach combined with *ex vivo* electrophysiological recordings. Sham and SDV mice received stereotaxic injections of a viral vector mixture (AAV-PPTA-Cre + AAV-FLEX-tdTomato or AAV-PPE-Cre + AAV-FLEX-tdTomato) into the NAc or DS (**Fig. 7A**). This strategy ^32^ enables the selective sparse labelling (**Fig. 7A**), visualization, and patch-clamp recording of D1R-expressing (PPTA, ^23^) and D2R-expressing (PPE, ^23^) SPNs (**Fig. 7B**). Using whole-cell patch-clamp electrophysiology, we investigated whether the gut-brain vagal axis, by modulating the mesolimbic DA system activity, influences the passive and active membrane properties of D1R- and D2R-SPNs in the NAc and DS. Interestingly, our recordings revealed an increased excitability of NAc D2R-SPNs in SDV mice, characterized by elevated membrane resistance (Ri), reduced rheobase, increased action potential rise/decay ratio and enhanced Input/Output gain function (also illustrated by a higher firing rate at +50 pA) (**Fig. 7C, C^1^**). Notably, no substantial differences were observed in NAc D1R-SPNs between experimental groups for all electrophysiological parameters (**Fig. 7C, C^1^**; see also **Suppl. Table 2**); apart from a modest but significant increase in firing rate at +50 pA in SDV mice which did not reflect in the overall Input/Output gain function. Interestingly, no significant changes were detected in DS D1R- or D2R-SPNs across experimental groups (**Fig. 7D, D^1^**; see also **Suppl. Table 3**).

**Figure 7:**
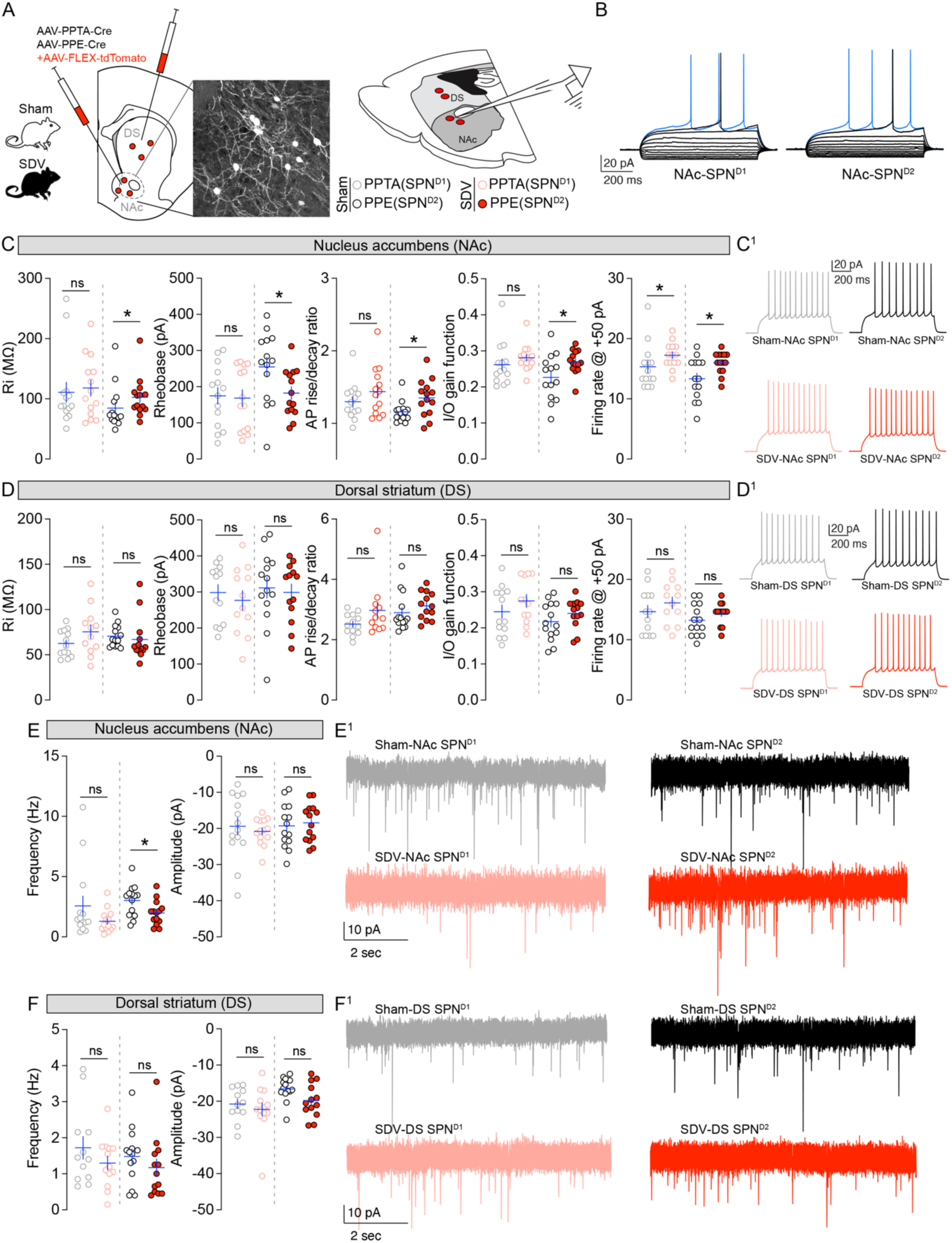
Gut-brain vagal inputs differentially scale the neuronal properties of D1R- and D2R-SPNs. (**A**) Drawing illustrates the double viral strategy to differentiate between D1R (PPTA)- and D2R (PPE)-spiny projection neurons (SPNs) in the NAc and DS. Inset shows immunofluorescence of sparse neuronal labeling. (**B**) Classical spiking profile of D1R- and D2R-SPNs. (**C-D^1^**) Passive and active membrane properties (membrane resistance; rheobase; action potential of rise/decay ratio; input/output grain function; firing rate at +50 pA) and spiking profiles of D1R- and D2R-SPNs in the NAc (**C, C^1^**) and DS (**D, D^1^**) in Sham (n=13-14 neurons) and SDV (n=12-14 neurons) mice. (**C-F^1^**) Frequency and amplitudes postsynaptic currents (PSCs) impinging onto D1R- and D2R-SPNs and their representative traces in the NAc (**E, E^1^**) and DS (**F, F^1^**) in Sham (n=12-14 neurons) and SDV (n=13-14 neurons) mice. Statistics: *p<0.05 for specific comparisons. Student’s t-test (C, D, E, F).

The increased excitability of NAc D2R-SPNs in SDV mice supports the hypothesis that alterations in DA release dynamics may drive these adaptations, given that ambient DA has a higher affinity for G_i_-coupled D2R. Moreover, we observed a lower frequency but equal amplitude of postsynaptic currents (PSCs) impinging onto NAc D2R-SPNs but not NAc D1R-SPNs (**Fig. 7E, E^1^**). Again, PSCs changes were restricted to NAc D2-SPNs since no differences were observed for DS D1R- and D2R-SPNs (**Fig. 7F, F^1^**).

These results highlight that the gut-brain vagal axis, by governing VTA DA-neuron activity and DA dynamics, differentially influence the responsiveness of NAc SPNs.

### The vagal tone contributes to orchestrating cell type-specific dendritic spines density in the NAc

Since DA plays a crucial role in regulating the dynamic morphological remodeling of SPNs, and given our findings suggesting that the gut-brain vagal tone influences mesolimbic DA dynamics and NAc SPNs activity, we investigated whether the structural morphology and density of dendritic spines in the NAc and DS depended on the integrity of the vagal pathway. To address this, we performed unbiased 3D diolistic reconstructions of dendritic spines in the NAc and DS of Sham and SDV mice. Our analysis revealed a significant reduction in dendritic spine density of NAc SPNs in SDV mice, whereas DS SPNs remained unaffected compared to control animals. This reduction was more pronounced in mushroom-like spines than in thin-like spines (**Fig. 8A-D**).

**Figure 8:**
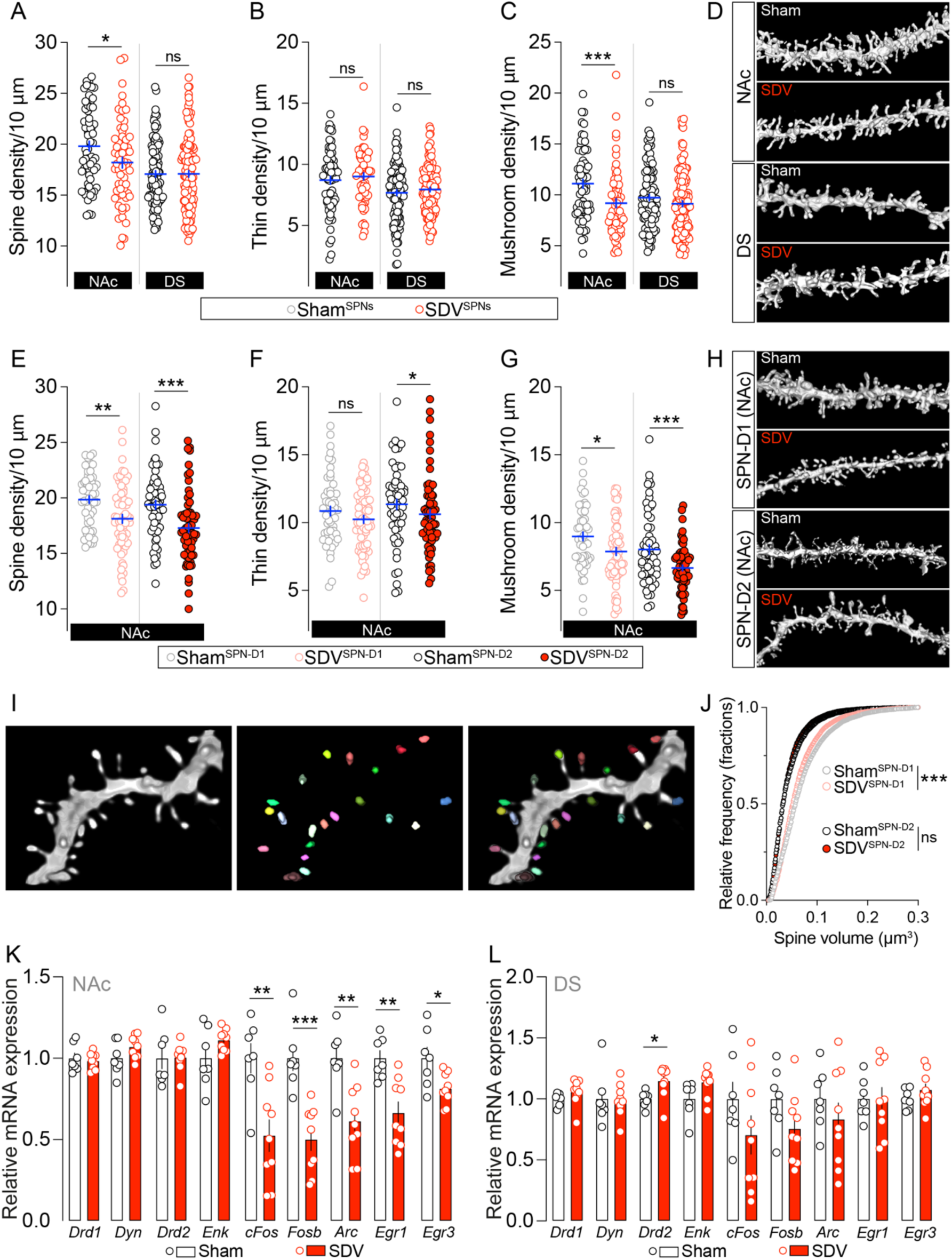
The integrity of the gut-brain vagal axis is necessary for the structural organization of dendritic spines in D1R- and D2R-SPNs. (**A**) Unbiased labeling of dendritic spine density and their classification [thin (**B**) and mushroom types (**C**)] in the NAc and DS of Sham (n=66-100 dendrites for NAc and DS groups) and SDV (n=60-127 dendrites for NAc and DS groups) mice. (**D**) 3D rendering of dendritic spines in the NAc and DS of Sham and SDV mice. (**E**) Viral cell type-specific labeling of dendritic spine density and their classification [thin (**F**) and mushroom types (**G**)] in the NAc Sham (n=53-59 dendrites for D1R- and D2R-SPNs groups) and SDV (n=59-65 dendrites for D1R- and D2R-SPNs groups) mice. (**H**) 3D rendering of dendritic spines in the NAc of Sham and SDV mice. (**I-J**) 3D rendering of a dendritic segment and the segmented spine heads. Segmentation procedure allows the measurement of head volumes from spines in NAc D1R- and D2R-SPNs of Sham and SDV mice. (**K, L**) Relative mRNA expression of SPNs markers (*Drd1*, *Dyn*, *Drd2*, *Enk*) and plasticity-related genes (*cFos*, *Fosb*, *Arc*, *Egr1*, *Egr3*) in the NAc and DS of Sham (n=6-7 for NAc and DS groups) and SDV (n=9 for NAc and DS groups) mice. Statistics: *p<0.05, **p<0.01 and ***p<0.001 for specific comparisons. Student’s t-test (A, B, C, E, F, G, J, K, L).

Given the heterogeneity of SPNs (D1R- *vs* D2R-expressing SPNs), we further examined potential cell type-specific differences using our viral sparse labelling strategy (**Fig. 7A**). Our results showed a marked decrease of spine density in both NAc D1R- and D2R-SPNs of SDV mice (**Fig. 8E, H**). However, distinct morphological adaptations emerged when analyzing dendritic spine subtypes (thin *vs* mushroom) and spine head volume. Specifically, NAc D2R-SPNs of SDV mice exhibited a reduction in both thin and mushroom spine densities, whereas in NAc D1R-SPNs, only mushroom spine density was significantly reduced (**Fig. 8F, G**). Furthermore, a pronounced decrease in the spine head volume was observed exclusively in NAc D1R-SPNs of SDV mice (**Fig. 8I, J**).

These findings indicate that dopaminoceptive Nac SPNs undergo structural remodeling following the disruption of the interoceptive vagal relay, a process likely mediated by vagal tone-dependent regulation of VTA DA-projecting neurons.

These structural modifications were not associated with significant alterations in key markers of D1R- and D2R-SPN identity, such as *Drd1*, *Dyn*, *Drd2*, and *Enk* (**Fig. 8K**), except for a slight increase in *Drd2* expression in the DS (**Fig. 8L**). However, we observed structure-specific changes in the expression of activity/plasticity-dependent immediate early genes (*cFos*, *Fosb*, *Arc*, and *Egr1*), which were altered in the Nac but not in the DS (**Fig. 8K, L**).

These latter findings suggest that vagus nerve-associated DA dynamics (**Fig. 3, 4**), electrophysiological changes of VTA DA-neurons (**Fig. 5**) as well as molecular, electrophysiological, and morphological adaptations of Nac SPNs (**Fig. 6, 7, 8**) may collectively shape the intrinsic regulation of plasticity markers in response to the disruption of vagal interoceptive inputs and therefore constitute a functional base for vagus-mediated reinforced behaviors (**Fig. 1, 2**).

## Discussion

Reward events, whether elicited by food or drugs of abuse, and their ensuing (mal)adaptive consequences, depend on the precise orchestration of distinct neural circuits and ensembles. These systems, which differentially process natural stimuli *versus* drugs of abuse, yet converge on modulating the activity of DA-neurons and DA-dependent dynamics. To date, much of our understanding has been shaped by an exteroceptive conceptual framework focused on external stimuli and environmental cues. However, recent insights are expanding this perspective by highlighting the crucial role of interoception, the unconscious sensing of the body’s internal physiological states, as a fundamental biological and mechanistic variable in the regulation of molecular, cellular, circuit-level, and behavioral responses.

In this study, by leveraging complementary, multi-scale, and integrative approaches, we identify the gut-brain vagal axis as an “essential modulator and extended component” of the DA reward system. Our findings reveal that the integrity of this axis is not merely supportive but plays a permissive and essential role in gating both the constitutive/spontaneous and stimulus-evoked activity of the mesolimbic DA circuit. Critically, we show that this modulation extends to the rewarding, reinforcing and conditioning properties of both natural rewards (*e.g.*, palatable food) and drugs of abuse as demonstrated by the use of a large panel of behavioral tests. These novel insights expand upon and deepen the implications of recent seminal studies mainly focused on food-related vagal signaling ^6,7,15,54^, contributing to the formulation of a more unified and holistic framework for understanding reward dynamics that integrates both interoceptive and exteroceptive domains. Specifically, our findings provide compelling evidence that the gut-brain vagal axis, via an indirect and polysynaptic periphery-to-brain pathway ^6^, exerts functional control over the activity and *in vivo* DA release dynamics of VTA DA-neurons, particularly within the mesolimbic VTA→Nac circuit. Disruption of this axis led to attenuated molecular responses, such as reduced expression of immediate early genes (cFos, Egr1), and diminished *in vivo* burst firing of VTA DA-neurons, which was associated with a lower probability of stimulus-evoked DA release events. Notably, the concentrations of dopamine, other bio amines (NE, 5-HT), and their metabolites, as well as markers of DA synthesis and metabolism, remained unaltered in SDV mice. This suggests that the vagal tone predominantly modulates the activity-dependent state of VTA DA-neurons, rather than their capacity for DA synthesis per se. Consistent with this, *ex vivo* recordings revealed that this blunted activity likely results from a reduced glutamatergic drive onto DA-neurons, accompanied by decreased expression of key glutamate-related transcripts in the VTA (*Grin1*, *Gria1*, *Grm1*). These findings align with previous work showing that *Grin1*/NR1-dependent bursting of VTA DA-neurons is essential for post-ingestive food-seeking behaviors ^7^. Whether this glutamatergic input is embedded within the heterogeneous VTA microcircuit, comprising Glu and dual Glu/DA-neurons ^55^, or arises from upstream glutamatergic projections remains to be elucidated. Of note, recent studies have identified two brainstem structures associated with vagal signaling that project to the VTA: the nucleus tractus solitarius (NTS, ^56^), the primary recipient of vagal afferents, and the parabrachial nucleus (PBN, ^57–59^). However, while we observed strong PBN→VTA connections, our tracing results revealed only a few and scattered NTS-neurons projecting to the VTA, suggesting the involvement of a vagus→NTS→PBN→VTA circuit, similarly to the previously reported vagus→NTS→PBN→SNpc circuit ^6^. It is therefore reasonable to hypothesize that chronic perturbation of the interoceptive gut-brain vagal relay, as exemplified by our mouse model or by pathological states (*i.e.*, obesity ^16^, degenerative disorders ^60–62^), may dampen the activity of the NTS→PBN→VTA circuit and thus consequently alters the mesolimbic VTA→NAc reward DA system. In fact, we noticed that this vagus nerve-mediated modulation profoundly shaped the molecular, cellular, morphological, and functional underpinnings of DA-dependent motivational and reinforced behaviors. Notably, our work extends previous observations linking vagal inputs to DA activity in the nigrostriatal SNpc→DS circuit ^6^, demonstrating that palatable food-evoked DA responses, either elicited by orosensory or intragastric stimuli, also robustly engage the mesolimbic VTA→NAc system. These results mechanistically reinforce a previous study showing that post-ingestive signals activate DA dynamics in humans ^63^. Furthermore, we show that the gut-brain vagal axis modulates DA dynamics and reward-related behaviors in the NAc, but not in the DS, in response to drugs of abuse, highlighting its specific relevance to mesolimbic, but not nigrostriatal, DA signaling. It is important to highlight that our work aligns with recent human studies showing that transcutaneous auricular vagus nerve stimulation (taVNS) increased the drive to work for non-food rewards ^64^ and promoted the activity of the DA midbrain ^65^.

In addition to changes in the DA midbrain, disruption of vagal integrity was associated with both electrophysiological and morphological alterations in the two major subtypes of NAc spiny projection neurons (SPNs), D1R- and D2R-expressing SPNs, but not in their DS counterparts. Specifically, disruption of the gut-brain vagal path led to reduced spine density in both NAc D1R- and D2R-SPNs reflecting structural remodeling likely driven by attenuated VTA DA release. While spine classification (mushroom *versus* thin) and spine head volume exhibited some subtype-specific variations, the overall trend pointed to synaptic destabilization of both cell types. These morphological results mirror other studies showing the pivotal role of DA in dendritic spine growth ^66–69^. Functionally, the most pronounced electrophysiological alterations occurred within NAc D2R-SPNs, which displayed heightened excitability across several passive and active membrane properties. This increased excitability may stem from reduced phasic/tonic activity of VTA DA-neurons, leading to decreased DA release and disinhibition of D2R-SPNs, given the inhibitory nature of Gi-coupled D2 receptors ^70^. Importantly, this supports the notion that changes in DA dynamics, beyond the traditional receptor affinity model ^71^, can differentially affect D1R and D2R signaling ^72,73^. In line with these changes, our findings also demonstrate that vagal integrity is essential for driving D1R- and D2R-mediated behaviors, as well as for the activity-dependent recruitment (cFos expression) of both NAc D1R- and D2R-SPNs, as confirmed using receptor-specific pharmacological tools. Here, we focused on the two major dopaminoceptive cell types in the DS/NAc. However, these territories also contain diverse interneuron populations, some of them also DA-sensitive (*i.e.*, cholinergic D2R/ChAT-interneurons), that play a crucial role in modulating SPN activity ^74^. Whether vagus-mediated regulation of VTA DA-neurons also alters the functional dynamics of local interneuron-to-SPN microcircuits remains to be determined. Indeed, a key limitation of our study lies in the use of the SDV model which was chosen to chronically and unbiasedly (no specific topography or cell types) perturbates the gut-brain interoceptive relay. However, while selective for gut-brain, rather than general periphery-brain, signaling, this model fails to account for the remarkable cellular heterogeneity of vagal sensory neurons. These neurons, although glutamatergic (VGLUT2-expressing neurons), comprise at least 24 distinct subtypes ^75^, implicated in transmitting and regulating internal physiological states including cardiac, immune, respiratory, feeding, and digestive functions ^18,33,42,76,77^. Despite technical and microsurgical challenges associated with gut and nodose manipulations in mouse models, future efforts should prioritize the use of more refined viral and transgenic strategies to selectively target gut-to-brain vagal subpopulations and clarify their distinct and specific roles in modulating mesolimbic DA dynamics in response to both natural and recreational/psychoactive rewards. Although technologies to selectively inhibit the vagus nerve in humans are not yet available, such refined approaches could help disentangle the distinct contributions of vagal subcircuits to specific aspects of reward-related behavior, thereby paving the way for new therapeutic strategies.

In conclusion, our findings position the gut-brain vagal axis as a fundamental interoceptive regulator of mesolimbic dopamine signaling, revealing its essential role in scaling the neural, molecular, and behavioral architecture of reward events and underscoring its potential as a therapeutic target for disorders marked by dysregulated motivation, reinforcement, or addiction. More broadly, our study contributes to a conceptual shift toward an expanded framework in which gut-derived interoceptive signals are recognized as integral components of reward processing, complementing and modulating traditionally brain-centric models of motivation, reinforcement and even drug addiction.

## Acknowledgments

We thank Olja Kacanski for administrative support; Isabelle Le Parco, Daniel Quintas, Magguy Boa, Ludovic Maingault, Angélique Dauvin and Florianne Michel for animals’ care. We also thank Dr. Enrica Montalban, Dr. Vincent Paillé, Charles Le Ciclé and Valerio Anelli for help, technical and experimental advice. We acknowledge the technical platform Functional and Physiological Exploration platform (FPE) of the Université Paris Cité, CNRS, *Unité de Biologie Fonctionnelle et Adaptative*, and the animal core facility “Buffon” of the Université Paris Cité/Institut Jacques Monod. We thank Fabrice Licata and the microscopy facility of the Campus Saint Germain des Prés (CSGP). We thank the Bioprofiler platform (Unit “*Biologie Fonctionnelle et Adaptative*”, Université Paris Cité, BFA, UMR 8251 CNRS, F-75205 Paris, France) for providing HPLC facilities and Justine Renault for technical assistance.

## Author contributions

O.O. initiated and developed the project, performed and analyzed most of the experiments, critically contributed to the overall design and execution. F.A. performed and analyzed 3D morphological experiments. T.L.B. performed and analyzed *in vivo* and *ex vivo* electrophysiology (VTA). S.P. performed and analyzed *ex vivo* electrophysiology on (DS and NAc). J.C. performed surgeries and animal caring. A.A, B.B., N.M, C.d.A. contributed to performing experiments. L.C.B. performed HPLC analyses. M.V. provided expertise in data analysis. S.L. provided critical feedback to the manuscript. L.V. supervised electrophysiology experiments (DS and NAc) and provided critical feedback to the manuscript. N.H. supervised 3D morphological experiments, provided critical feedback to the manuscript, contributed to experimental plan and funding. F.M. supervised electrophysiology experiments (VTA), provided critical feedback to the manuscript, contributed to experimental plan and funding. G.G. supervised the whole project, secured funding, provided guidance, designed the experimental plan, wrote and finalized the manuscript with the help of the co-authors.

## Data availability

All the data generated have been included in the article. Datasets are available from the corresponding author upon reasonable request.

---

**Suppl. Figure 1:**
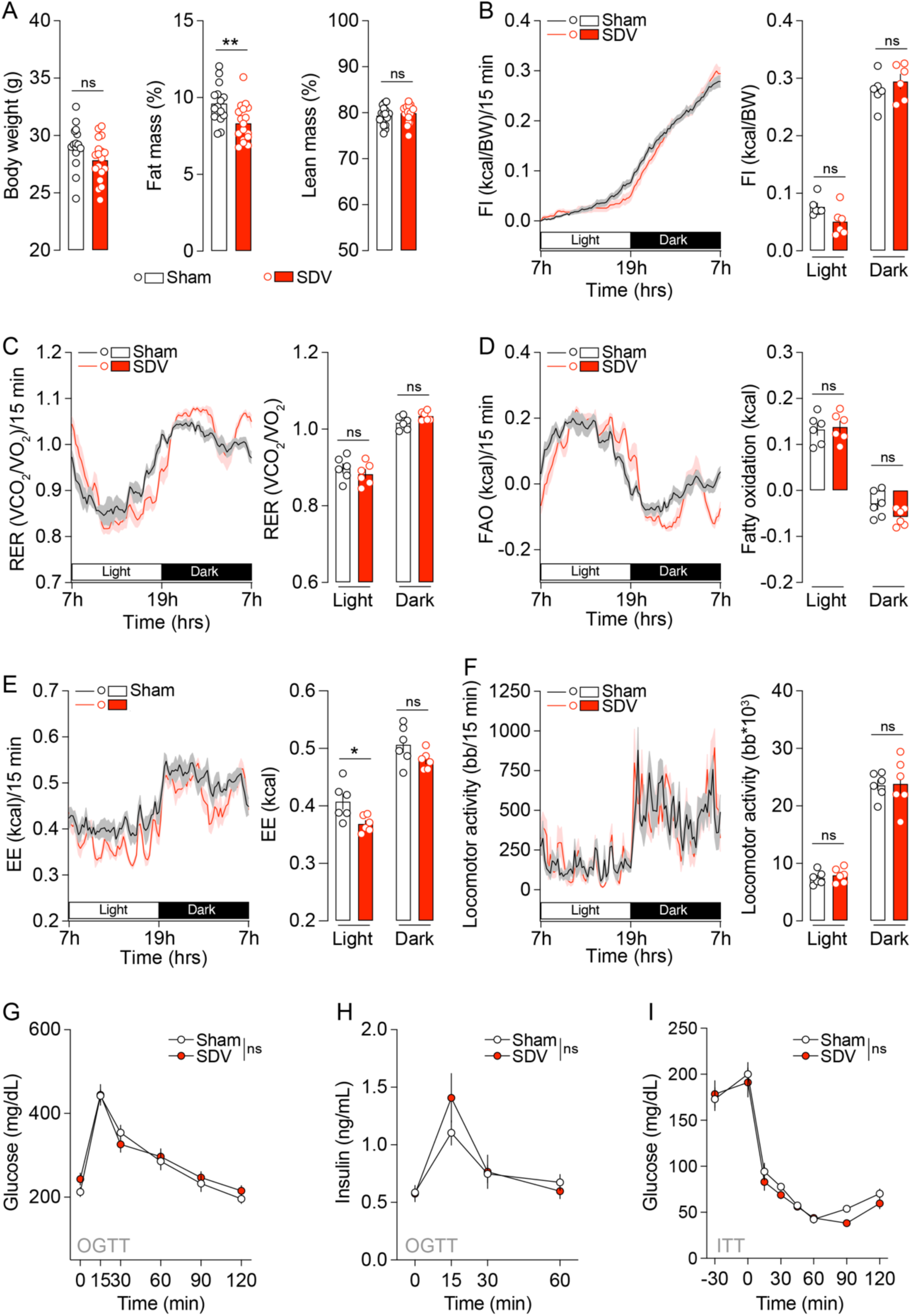
Metabolic profile of Sham and SDV mice. (**A**) Body weight, fat and lean mass of Sham (n=15) and SDV (n=18) mice. (**B-F**) Measurements of food intake (FI), respiratory exchange ratio (RER), fatty acid oxidation (FAO), energy expenditure (EE) and locomotor activity in Sham (n=6) and SDV (n=6) mice. (**G-H**) Glucose and insulin profiles following an oral glucose tolerance test (OGTT) in Sham (n=10) and SDV (n=9) mice. (**I**) Glucose profile following an insulin tolerance test (ITT) in Sham (n=9) and SDV (n=8) mice. Statistics: *p<0.05 for specific comparisons. Student’s t-test (A); Two-way ANOVA (B, C, D, E, F, G, H, I).

**Suppl. Figure 2:**
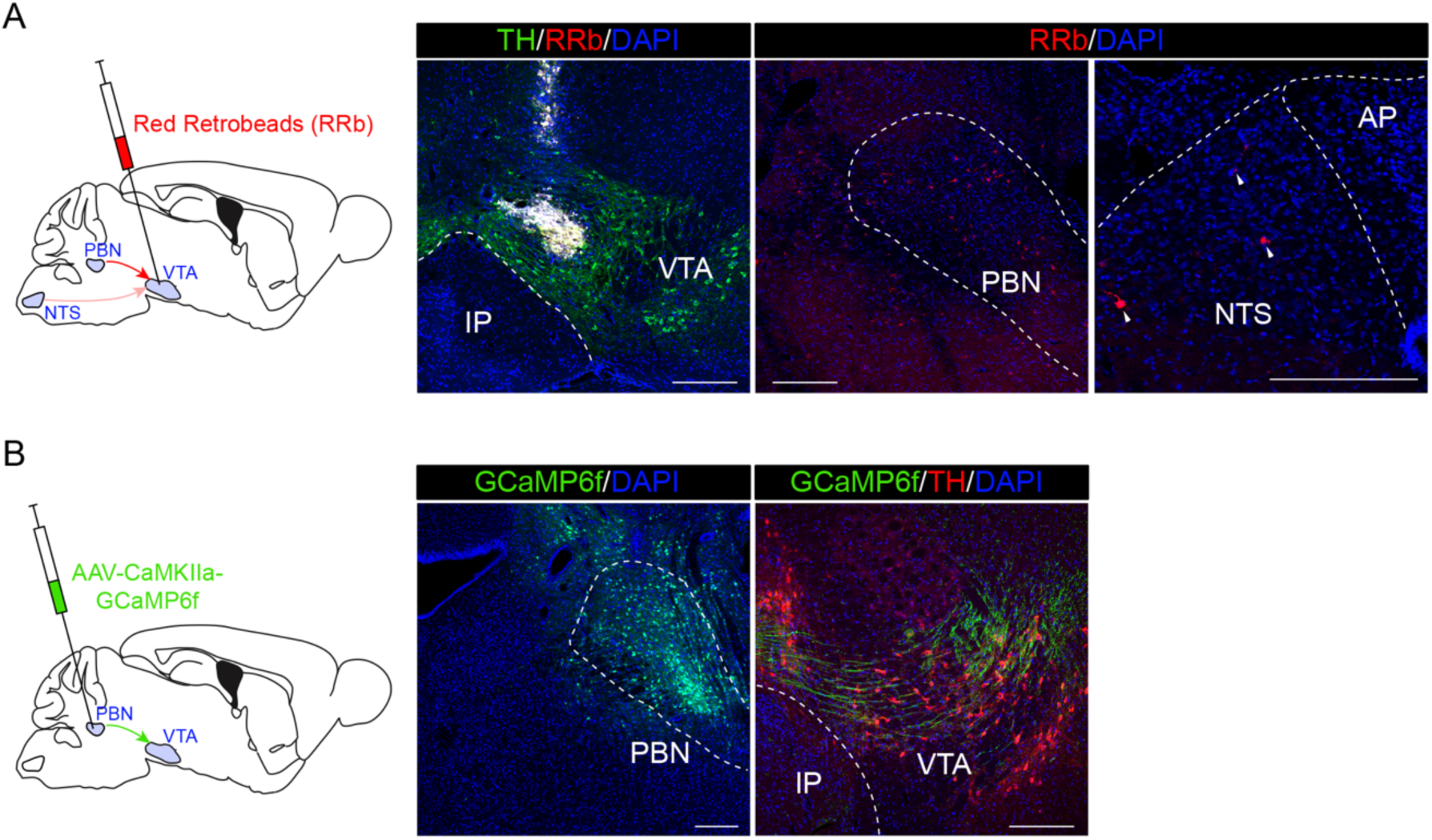
Connections from brainstem nuclei to the VTA. (**A**) Drawing illustrates the retrograde tracing approach [Red Retrobeads (RRb)]. Images show the injection site (VTA) and RRb-positive neurons in the PBN and NTS. (**B**) Drawing illustrates the anterograde viral (AAV-CaMKIIa-GCaMP6f) strategy. Images show the injection site (PBN) and GCaMP6f-positive terminals in the VTA. Scale bars: 250 μm.

**Suppl. Figure 3:**
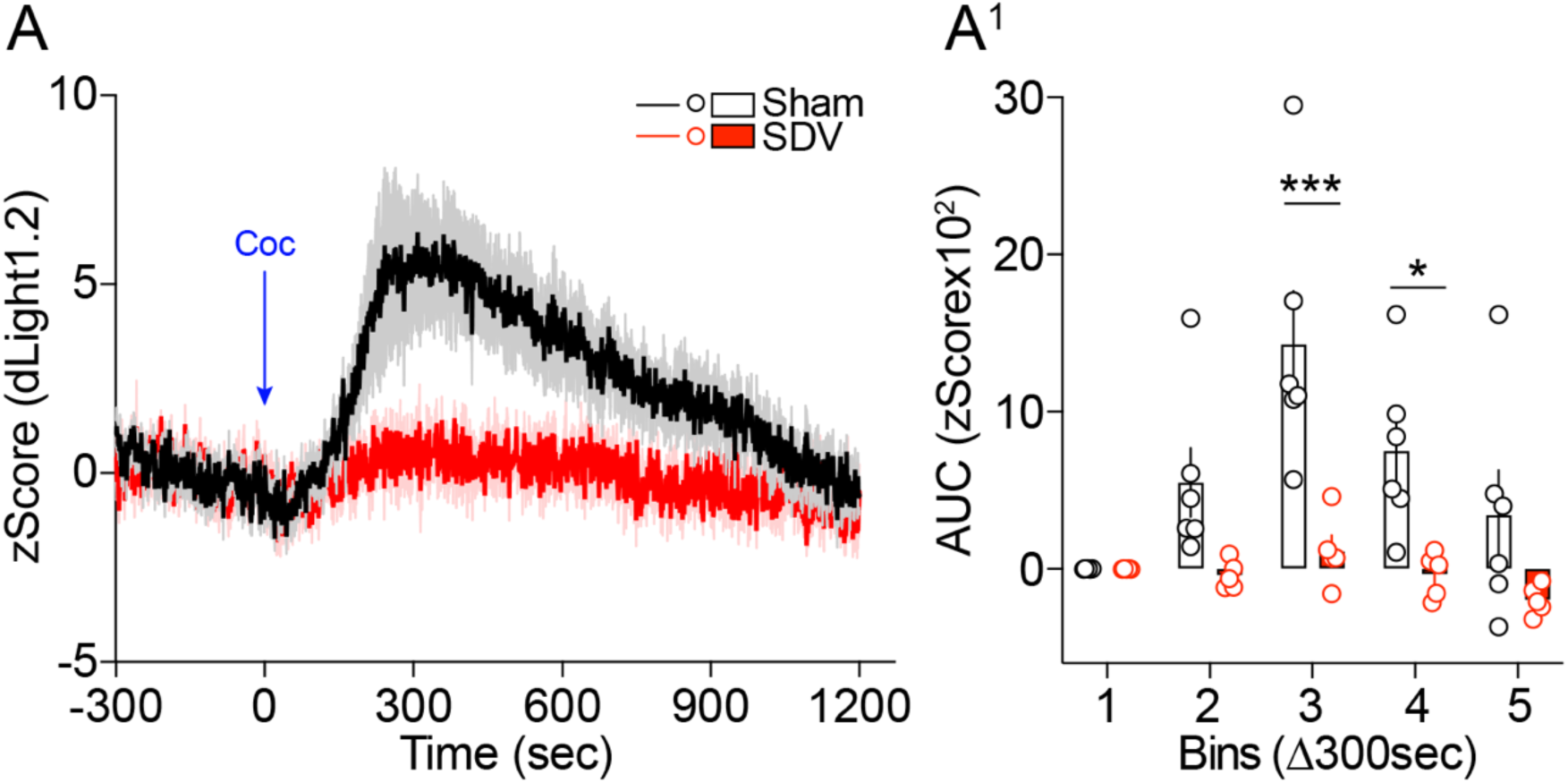
*In vivo* DA transients following administration of cocaine. (**A, A^1^**) *In vivo* DA dynamics during cocaine-induced DA release/accumulation in the NAc of Sham (n=6) and SDV (n=5) mice using the dLight1.2 biosensor. Statistics: *p<0.05 and ***p<0.001 for specific comparisons. Two-way ANOVA (A^1^).

**Suppl. Figure 4:**
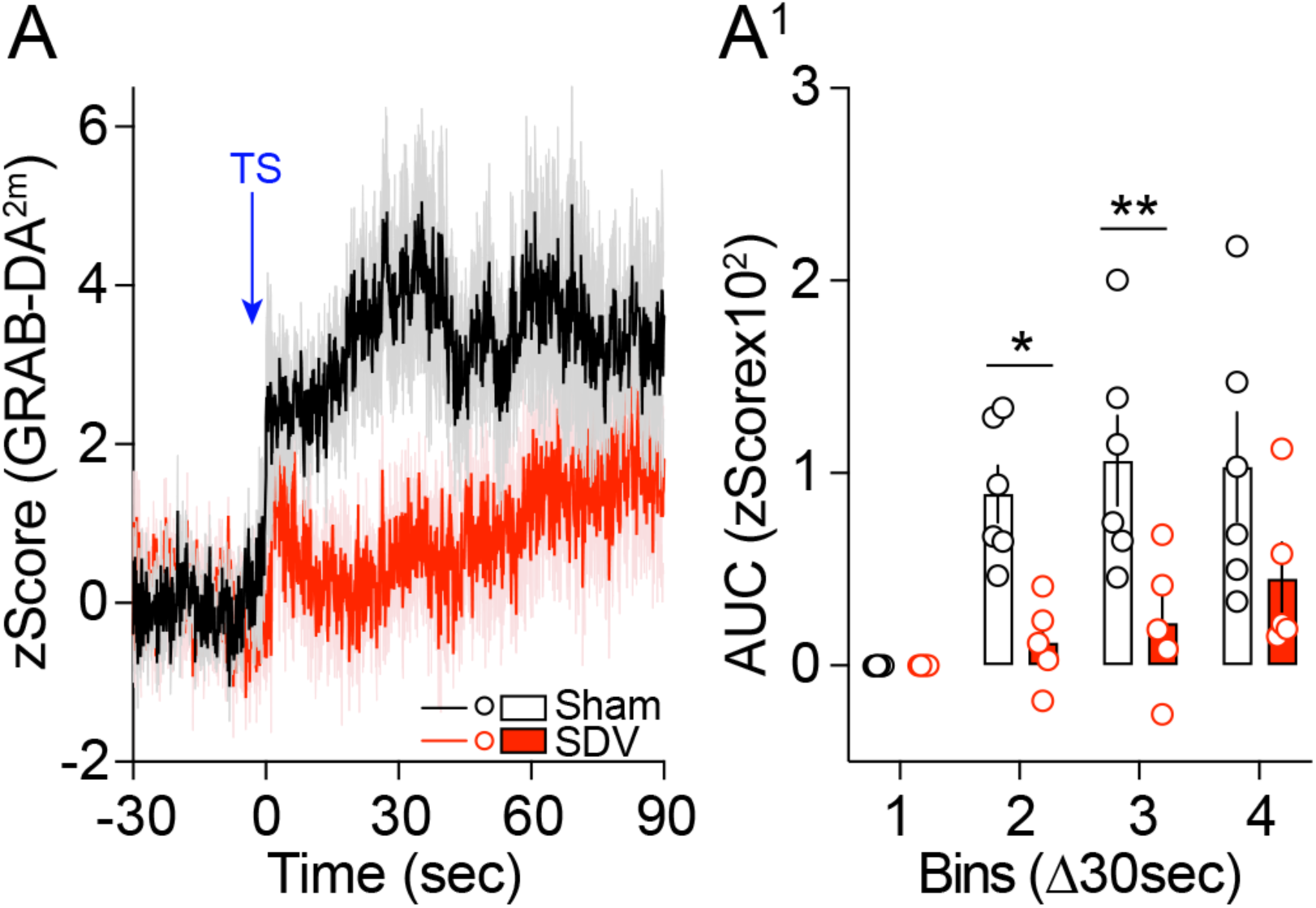
*In vivo* DA transients following a tail suspension test. (**A, A^1^**) *In vivo* DA dynamics during a tail suspension (TS)-induced DA release/accumulation in the NAc of Sham (n=6) and SDV (n=5) mice. Statistics: *p<0.05 and **p<0.01 for specific comparisons. Two-way ANOVA (A^1^).

**Suppl. Figure 5:**
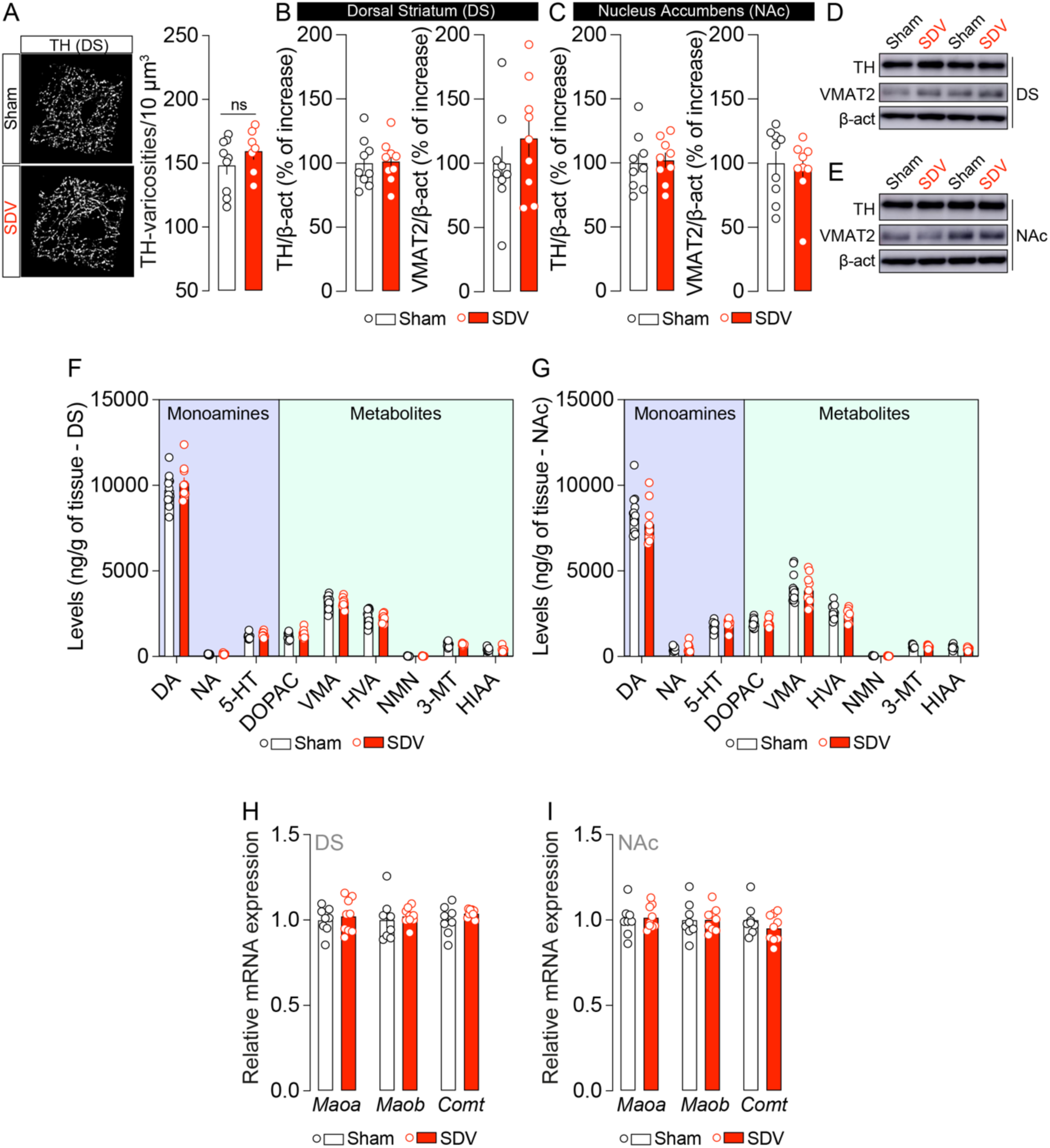
The integrity of the gut-brain vagal axis is not necessary for DA synthesis and metabolism. (**A**) 3D reconstruction and computational segmentation of VTA-projecting TH-positive varicosities in the NAc of Sham (n=9) and SDV (n=7) mice. (**B-E**) Expression and representative blots of TH and VMAT2 in the DS (**B, D**) and NAc (**C, E**) of Sham (n=9) and SDV (n=9) mice. (**F, G**) Quantifications of monoamines [dopamine (DA), noradrenaline (NA), serotonin (5-HT)] and their metabolites (DOPAC, VMA, HVA, NMN, 3-MT and HIAA) in the DS and NAc of Sham (n=12) and SDV (n=10) mice. (**H, I**) Relative expression of *Maoa*, *Maob*, Comt in the DS and NAc of Sham (n=8) and SDV (n=9) mice. Statistics: Student’s t-test (A, B, C, F, G, H, I).

**Suppl. Table 1.**
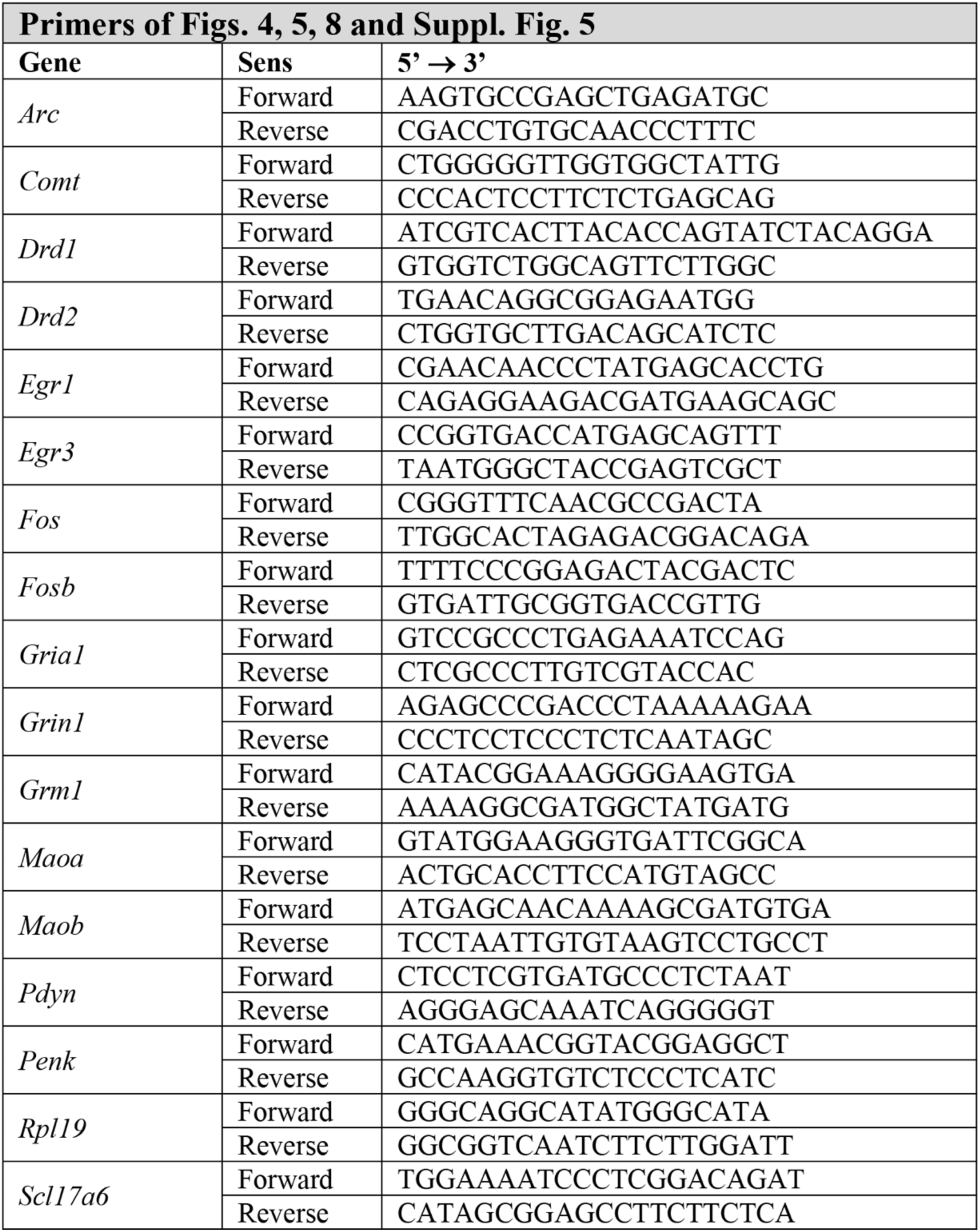

**Suppl. Table 2.**

| 1 | Nucleus accumbens (NAc) |  |  |  |  |  |
| --- | --- | --- | --- | --- | --- | --- |
|  | D1-SPN |  |  | D2-SPN |  |  |
|  | Sham<br>(n=14) | SDV<br>(n=14) | Stats<br>(p value) | Sham<br>(n=14) | SDV<br>(n=14) | Stats<br>(p value) |
| **Passive membrane properties** |  |  |  |  |  |  |
| RMP (mV) | -75.86 ± 1.565 | -78.64 ± 0.992 | 0.1155 | -79.50 ± 0.789 | -77.57 ± 1.357 | 0.0868 |
| Ri (MOhms) | 111.0 ± 16.70 | 118.1 ± 50.16 | 0.5714 | 84.54 ± 9.724 | 102.6 ± 8.763 | * 0.0274 |
| Membrane time constant (ms) | 7.953 ± 1.495 | 7.416 ± 1.436 | 0.8036 | 6.047 ± 0.6363 | 6.657 ± 0.6985 | 0.4824 |
| Inward rectification ratio | 0.7964 ± 0.0280 | 0.7810 ± 0.0338 | 0.9820 | 0.8461 ± 0.0251 | 0.8009 ± 0.0224 | 0.2100 |
| **Action potential properties** |  |  |  |  |  |  |
| Rheobase | 174.7 ± 22.85 | 168.4 ± 23.15 | 0.8132 | 254.9 ± 26.48 | 182.4 ± 17.59 | * 0.0156 |
| AP threshold (pA) | -34.49 ± 2.232 | -35.60 ± 1.331 | >0.9999 | -35.00 ± 1.207 | -36.21 ± 0.7837 | 0.5714 |
| Delay to 1st spike (ms) | 510.9 ± 30.25 | 449.9 ± 29.08 | 0.1167 | 571.4 ± 27.11 | 507.4 ± 39.33 | 0.2852 |
| AP Amplitude (pA) | 80.32 ± 2.161 | 84.68 ± 2.620 | 0.0690 | 84.67 ± 1.453 | 82.99 ± 1.863 | 0.3519 |
| AP rise time (ms) | 0.4111 ± 0.0143 | 0.4332 ± 0.0253 | 0.4752 | 0.4086 ± 0.0204 | 0.4071 ± 0.0129 | 0.9729 |
| AP rise/decay ratio | 1.299 ± 0.0667 | 1.438 ± 0.0901 | 0.2700 | 1.159 ± 0.0384 | 1.351 ± 0.0677 | * 0.0359 |
| **Action potential trains** |  |  |  |  |  |  |
| I/O gain function | 0.2618 ± 0.0161 | 0.2811 ± 0.0096 | 0.0776 | 0.2264 ± 0.0170 | 0.2694 ± 0.0088 | * 0.0404 |
| Firing rate at +50 pA | 15.33 ± 0.986 | 17.24 ± 0.567 | * 0.0226 | 13.33 ± 0.937 | 16.00 ± 0.484 | * 0.0310 |
| SFA ratio | 1.272 ± 0.1957 | 1.156 ± 0.0944 | 0.9820 | 0.981 ± 0.0364 | 1.019 ± 0.0726 | 0.6260 |

**Suppl. Table 3.**

| 2 | Dorsal striatum (DS) |  |  |  |  |  |
| --- | --- | --- | --- | --- | --- | --- |
|  | D1-SPN |  |  | D2-SPN |  |  |
|  | Sham<br>(n=13) | SDV<br>(n=12) | Stats<br>(p value) | Sham<br>(n=14) | SDV<br>(n=13) | Stats<br>(p value) |
| **Passive membrane properties** |  |  |  |  |  |  |
| RMP (mV) | -80.08 ± 0.780 | -80.00 ± 0.674 | 0.8611 | -80.93 ± 1.495 | -79.54 ± 1.153 | 0.1338 |
| Ri (MOhms) | 62.55 ± 4.058 | 75.47 ± 7.525 | 0.1683 | 70.55 ± 3.298 | 66.96 ± 6.778 | 0.0850 |
| Membrane time constant (ms) | 5.086 ± 0.415 | 5.093 ± 0.518 | 0.8938 | 6.066 ± 0.530 | 6.703 ± 1.202 | 0.8300 |
| Inward rectification ratio | 0.8705 ± 0.0304 | 0.8816 ± 0.0276 | 0.9787 | 0.8280 ± 0.0225 | 0.8519 ± 0.0278 | 0.4583 |
| **Action potential properties** |  |  |  |  |  |  |
| Rheobase | 298.5 ± 22.14 | 276.7 ± 27.10 | 0.4785 | 311.6 ± 28.28 | 299.0 ± 22.79 | 0.6500 |
| AP threshold (pA) | -39.33 ± 1.250 | -38.72 ± 1.009 | 0.9362 | -37.63 ± 0.9604 | -40.27 ± 1.263 | 0.1021 |
| Delay to 1st spike (ms) | 501.7 ± 37.56 | 477.0 ± 39.84 | 0.5033 | 501.0 ± 32.58 | 495.9 ± 32.55 | 0.9810 |
| AP Amplitude (pA) | 93.96 ± 1.820 | 93.98 ± 2.783 | 0.9048 | 90.85 ± 1.761 | 96.45 ± 1.602 | * 0.0427 |
| AP rise time (ms) | 0.4023 ± 0.0110 | 0.3817 ± 0.0143 | 0.8202 | 0.3671 ± 0.0175 | 0.3890 ± 0.00850 | 0.6066 |
| AP rise/decay ratio | 2.516 ± 0.0969 | 2.970 ± 0.2613 | 0.1519 | 2.899 ± 0.1653 | 3.123 ± 0.1264 | 0.1159 |
| **Action potential trains** |  |  |  |  |  |  |
| I/O gain function | 0.2450 ± 0.0187 | 0.2746 ± 0.0179 | 0.2993 | 0.2171 ± 0.01407 | 0.2400 ± 0.01121 | 0.2836 |
| Firing rate at +50 pA | 14.67 ± 1.0350 | 16.11 ± 0.9773 | 0.2832 | 13.24 ± 0.7053 | 14.36 ± 0.5042 | 0.2468 |
| SFA ratio | 0.8981 ± 0.0548 | 0.8707 ± 0.0263 | 0.9056 | 0.8868 ± 0.0342 | 0.9351 ± 0.0331 | 0.2800 |

